# SR7 - a dual function antisense RNA from *Bacillus subtilis*

**DOI:** 10.1101/2020.05.15.097949

**Authors:** Inam Ul Haq, Peter Müller, Sabine Brantl

**Affiliations:** Friedrich-Schiller-Universität Jena, Matthias-Schleiden-Institut, AG Bakteriengenetik, Philosophenweg 12, D-07743 Jena, Germany

**Keywords:** dual-function antisense RNA, *Bacillus subtilis*, regulatory peptide, SR7, SR7P, enolase, RNase Y, RNA degradation, degradosome

## Abstract

Here, we describe SR7, a dual-function antisense RNA from the *Bacillus subtilis* chromosome. This RNA was earlier published as the SigB-dependent regulatory RNA S1136 and reported to reduce the amount of the small ribosomal subunit under ethanol stress. We found that the 5’ portion of SR7 encodes a small protein composed of 39 amino acids which we designated SR7P. It is translated from a 185 nt SigB-dependent mRNA under five different stress conditions and a longer SigB-independent RNA constitutively. Two- to three-fold higher amounts of SR7P were detected in *B. subtilis* cells exposed to salt, ethanol or heat stress. Co-elution experiments with SR7P_C-FLAG_ and Far-Western blotting demonstrated that SR7P interacts with the glycolytic enzyme enolase. Enolase is a scaffolding component of the *B. subtilis* degradosome where it interacts with RNase Y and phosphofructokinase PfkA. We found that SR7P increases the amount of RNase Y bound to enolase without affecting PfkA. RNA does not bridge the SR7P-enolase-RNase Y interaction. *In vitro*-degradation assays with the known RNase Y substrates *yitJ* and *rpsO* mRNA revealed enhanced enzymatic activity of enolase-bound RNase Y in the presence of SR7P. Northern blots showed a major effect of enolase and a minor effect of SR7P on the half-life of *rpsO* mRNA indicating a fine-tuning role of SR7P in RNA degradation. Moreover, SR7P impacts survival of *B. subtilis* under stress conditions. We suggest that the SR7P-dependent modification of the degradosome affects targets in different physiological pathways.

## Introduction

Although short open reading frames (sORFs) are present in all genomes, they have been often missed in annotations. Therefore, small proteins (sproteins) comprising less than 50 amino acids (aa) are understudied. Nevertheless, a number of small proteins involved in different pathways have been identified serendipitously and investigated in more detail (rev. in [1, 2, 3]). Only recently, some focused efforts were made to study whole peptidomes (rev. in [4]). Among the few peptides studied so far are type I toxins that are often integrated into the membrane (rev. in [5]), chaperones of nucleic acids and metals, membrane components, factors stabilizing or disrupting larger protein complexes and regulatory peptides.

Type I toxins encompass usually <50 aa and either induce pores in bacterial membranes, act as RNase or DNase or interfere with cell envelope biosynthesis without affecting the membrane potential (rev. in [5]), like *B. subtilis* BsrG [6]. To be inserted into the membrane, these peptides carry trans-membrane domains.

RNA chaperones like Hfq (60-100 aa) and CsrA (≈60 aa) belong to small proteins and have been studied in much detail [7,8,9,10]. In addition, in *B. subtilis*, the small proteins FbpA (59 aa), FbpB (53 aa) and FbpC (29 aa) were proposed to be potential RNA chaperones [11]. Among them, FbpB has been shown to be required for the action of the sRNA FsrA on certain target mRNAs [12], although no biochemical analyses have been performed so far to demonstrate its RNA binding activity. By contrast, other peptides bind metal ions, as e.g. *E. coli* MntS (42 aa) that binds manganese and delivers it to other proteins [13].

SpoVM (26 aa) from *B. subtilis* [14] whose NMR structure has been solved recently [15] is an example for an sprotein as membrane component. It recognizes – most probably as multimer – convex membrane curvature, acts as a cue for the deposition of the endospore coat and tethers the endospore coat to the developing forespore [16].

An example for an sprotein interacting with a regulatory protein in *B. subtilis* is Sda (46 aa) that inhibits KinA, the first kinase needed for activation of the key regulator of sporulation Spo0A [17]. An sprotein that disrupts a large complex is the 40 aa comprising *B. subtilis* MciZ [18]. It binds to the C-terminal polymerization interface of FtsZ, where it causes shortening of protofilaments and blocks the assembly of higher order FtsZ structures [19].

Small regulatory RNAs (sRNAs), the most important posttranscriptional regulators in all three kingdoms of life, can act by base-pairing or by protein-binding [20-23]. Chromosome-encoded sRNAs inhibit or activate translation or affect RNA stability, although the detailed mechanisms vary. The majority of base-pairing sRNAs do not comprise an ORF and are, therefore, not translated. However, a handful of trans-encoded sRNAs contain an ORF and have two functions, a base-pairing and a peptide-encoding. They have been designated dual-function sRNAs [24]. To date, only one dual-function sRNA from Gram-negative bacteria, SgrS [25], is known and has been intensively studied, whereas the other dual-function sRNAs are encoded in the genomes of Gram-positive bacteria: RNAIII (rev. in [26]) and psm-mec RNA from *Staphylococcus aureus* [27], Pel RNA from *Streptococcus pyogenes* [28] and SR1 from *Bacillus subtilis* [29-32]. SR1 encodes a 39 aa sprotein, SR1P, which interacts with the glyceraldehyde-3P-dehydrogenase A (GapA), thereby promoting the interaction of GapA with RNase J1 and increasing RNase J1 activity [33]. On a few other sRNAs, small ORFs have only been detected, but it has not been elucidated so far whether or not they are translated and what their biological functions are [24].

We chose *ncr2360* RNA that was originally published in 2010 [34] among 54 newly identified sRNAs from *B. subtilis* to investigate the function of its encoded peptide. In preliminary experiments, we could detect the RNA in Northern blots and show – using a translational *ncr2360*-*lacZ* fusion under control of the heterologous promoter pIII [35] – that the putative *ncr2360* SD sequence is recognized in *B. subtilis*. Here, we rename *ncr2360* as *sr7* and report that *sr7* is transcribed from a SigB-dependent promoter under five stress conditions. Its encoded peptide SR7P (39 aa) is synthesized in *B. subtilis* and additionally produced to a low extent from a longer RNA originating at a SigA-dependent promoter located 1.6 kb upstream. We demonstrate that SR7P directly interacts with the glycolytic enzyme enolase which moonlights as scaffolding component in the putative *B. subtilis* degradosome [36]. The SR7P-enolase interaction promotes binding of RNase Y to enolase. SR7P does not seem to interact with the other scaffolding component, phosphofructokinase PfkA. *In vitro* RNA degradation assays and Northern blotting using two known RNase Y substrates reveal a contribution of SR7P to RNase Y-dependent RNA degradation. In addition, SR7P affects cell survival under ethanol, acid and heat stress. Interestingly, in 2015, SR7 was published as SigB-dependent antisense RNA S1136, which is convergently transcribed to *rpsD* RNA and affects the amount of the small ribosomal subunit under ethanol stress [37]. However, it escaped the authors’ attention that S1136 contains an ORF which might code for a small protein. Based on their and our data, SR7 is the second dual-function regulatory RNA found in *Bacillus subtilis*. In contrast to the trans-encoded sRNA SR1 [30, 38], SR7 is a cis-encoded *bona-fide* antisense RNA.

## Results

### Two independent RNA species are detectable in the intergenic region between *tyrS* and *rpsD*

The *sr7* gene is located on the *B. subtilis* chromosome between the *tyrS* and *rpsD* genes and transcribed convergently to the *rpsD* gene (Fig. 1A). As Mars *et al*. had previously reported that S1136 – here renamed as SR7 – is transcribed under control of SigB and induced under ethanol stress [37], we mapped the transcriptional start site of SR7 under non-stress, ethanol and acid stress conditions. To this end, *B. subtilis* DB104 was cultivated in TY medium, either induced with HCl (pH 5.0) or with 4 % ethanol or not induced, RNA prepared, treated with DNase and subjected to primer extension. For comparison, both an isogenic *ΔsigB* strain and a strain where the SigB-dependent promoter upstream of *sr7* was knocked out [DB104(*Δp*_*sr7*_)] grown under nonstress and acid stress conditions were used (Fig. 1B). Without treatment, we observed a weak transcription initiation signal at the expected position 14 bp downstream of the −10 box of the SigB promoter and a second, much stronger signal far upstream, which most probably originated at the *tyrS* promoter (Fig. 1C). After HCl or ethanol stress, the SigB-dependent signal was about twofold or 50-fold stronger, respectively, compared to non-stress conditions. These signals were missing in RNA from the *ΔsigB* strain as well as from DB104(*Δp*_*sr7*_) confirming that they originate at the SigB-dependent promoter p_*sr7*_.

**Fig. 1.**
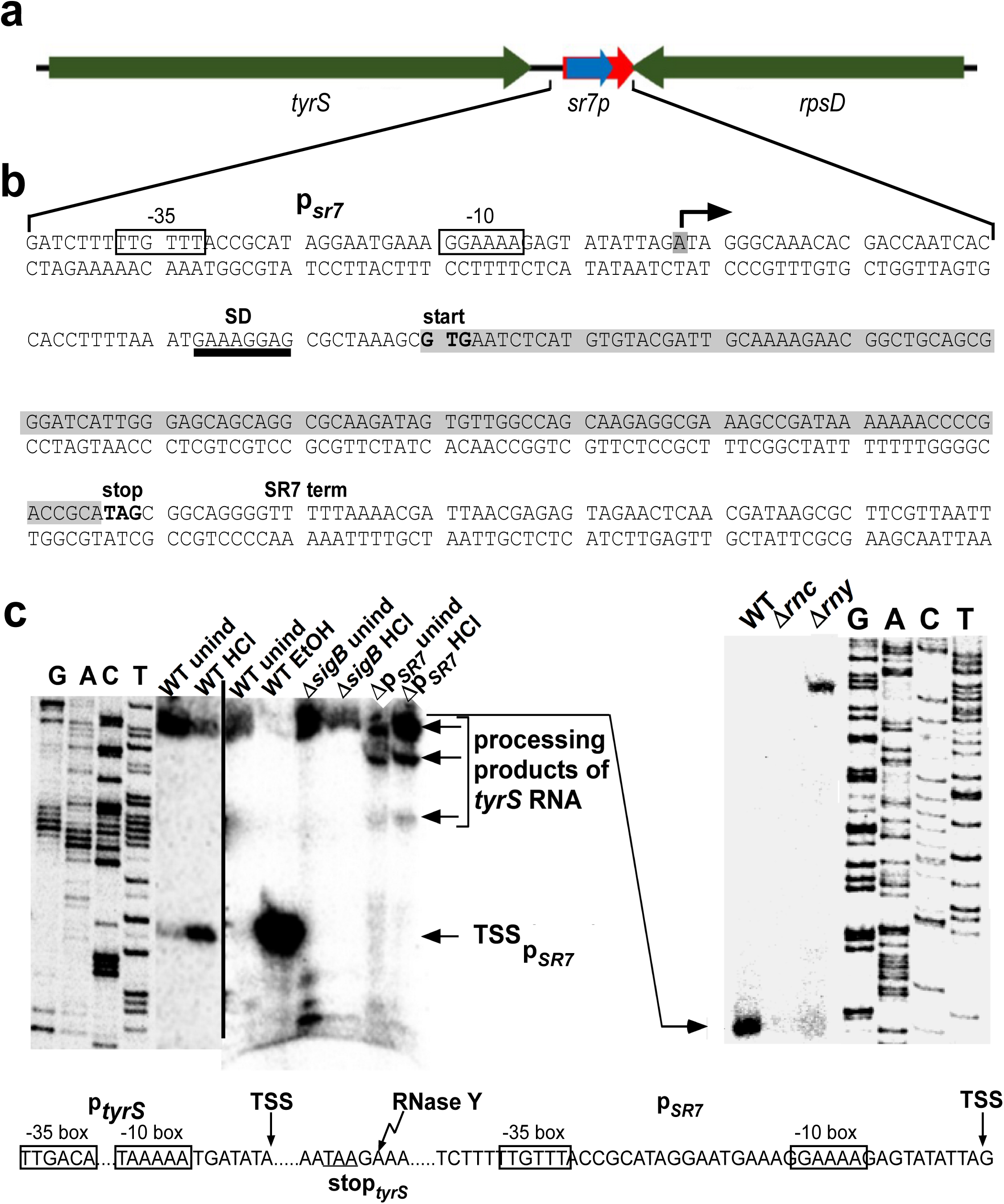
Location of the *sr7* gene and determination of transcription start and processing sites. (a) Schematic representation of the location of the *sr7* locus on the *B. subtilis* chromosome. The direction of transcription is indicated by arrows. (b) Sequence of the *sr7/rpsD* locus. −35 and −10 boxes of the *sr7* promoter is indicated. Start and direction of transcription are indicated by arrows and transcription termination is indicated by ‘*sr7* term’. Start and stop codons of the *sr7* ORF are in bold and the SD sequence is underlined. (c) Left: Mapping of the 5’ ends of SR7 under acid (HCl) and ethanol stress conditions. Primer extension was performed as described in *Materials and Methods*. Right: Mapping of the processing site of the *tyrS* promoter derived transcript in wild-type DB104 and isogenic *Δrnc* and *Δrny* strains. Below: Sequences of the *tyrS* and SigB-dependent promoters. TSS, transcription start site. Zigzag arrow, mapped RNase Y dependent processing site of the *tyrS* promoter derived transcript.

To find out if the bands upstream of the SR7 transcription start site were processing products or derived from an additional upstream promoter, we employed RNA from a *Δrny* and a *Δrnc* strain in comparison to the wild-type (right panel). In case of the *Δrny* strain, the signal was strongly reduced and instead a new band appeared 74 bp upstream, indicating that this band was a processing product from a longer RNA originating far upstream. As the only SigA-dependent promoter located upstream of p_*sr7*_ is the *tyrS* promoter, these signals most probably originated there. We mapped the processing site of the longer SR7 species directly 3’ of the *tyrS* stop codon (see Fig. 1C). In the case of the *Δrnc* strain, the original signal disappeared and no upstream signal was detectable. This suggests that RNase III might be either involved in cleavage of a duplex formed between transcripts originating at the *rpsD* promoter and those orginating at the *tyrS* promoter in the absence of stress or, alternatively, in cleavage of a longer double-strand formed within the *tyrS* transcript.

In summary, we conclude that two independent transcripts are observable in the intergenic *tyrS-rpsD* region, one – SR7 – originating at the SigB-dependent promoter p_*sr7*_ and a processing product of a transcript from the SigA-dependent *tyrS* promoter.

### The SigB-dependent p_sr7_ is induced under five different stress conditions

To analyse under which stress conditions *sr7* is transcribed from the SigB-dependent promoter, *B. subtilis* strain DB104 was grown in TY, either 4 % ethanol, 0.5 M NaCl, 1 mM manganese or HCl to pH 5.0 added or the culture shifted from 37 °C to 48 °C (heat shock). After 30 min stress, aliquots were flash-frozen, total RNA prepared and subjected to Northern blotting. Upon stress, a strong SR7 band of ≈185 nt appeared which was not visible under non-stress conditions (Figs. 2 and S1). No induction of SR7 was observed under oxygen stress or deficiency, vancomycin or iron stress (not shown). In a rifampicin experiment, a half-life of ≈29 min was determined for the 185 nt SR7 under NaCl stress conditions indicating that it is very stable (Fig. 2). The same stability was observed for SR7 after ethanol stress (Fig. S1).

**Fig. 2.**
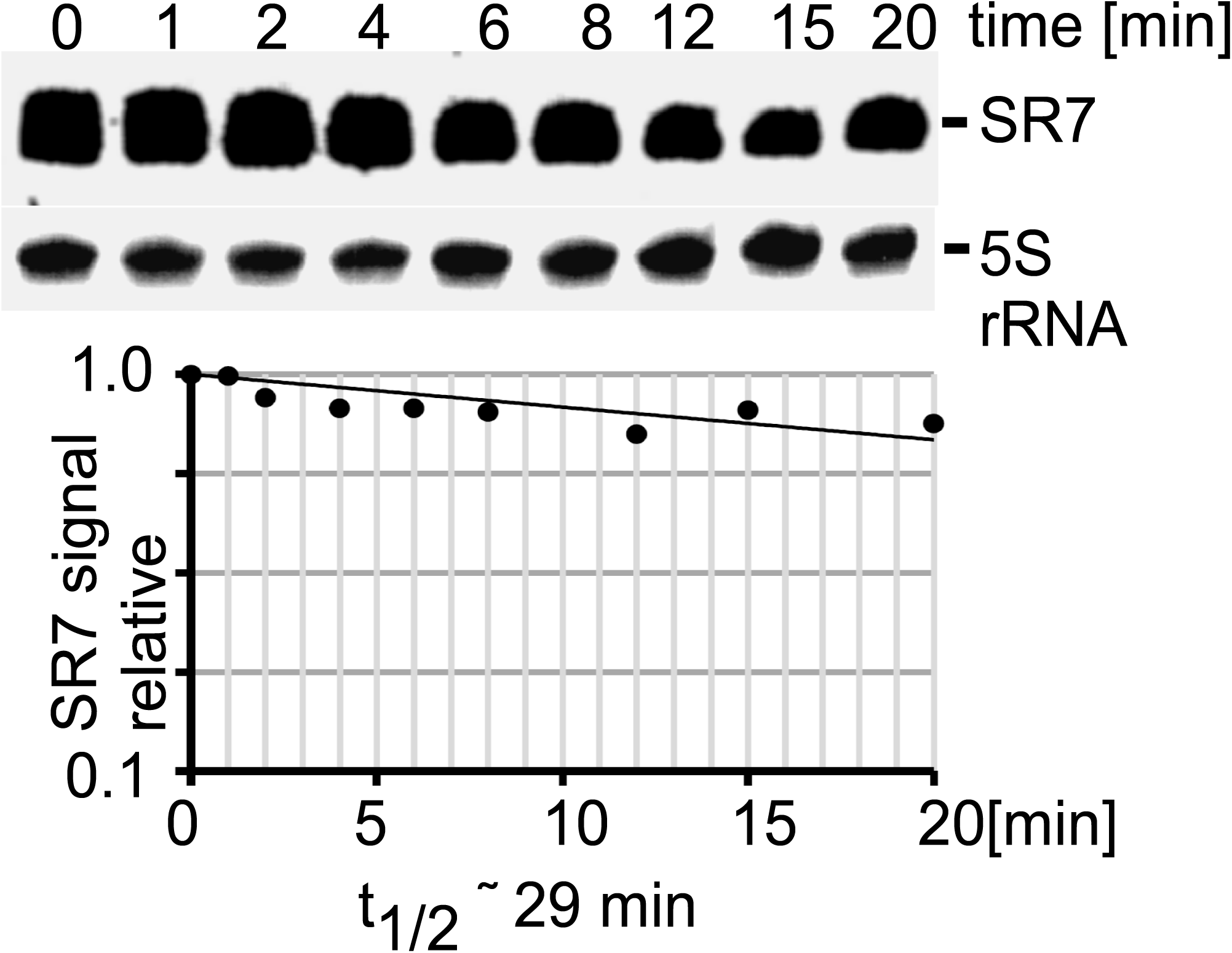
Determination of the half-life of SR7 under salt stress. *B. subtilis* DB104 was grown in TY medium until stationary phase at 37 °C, induced by 0.5 M NaCl for 15 min, then rifampicin added, time samples taken, total RNA prepared and subjected to Northern blotting. SR7 was detected by hybridization with a [^32^P]-αUTP-labelled riboprobe. The autoradiogram of the Northern blots is shown. Loading errors were corrected by reprobing with a [^32^P] γ-ATP-labelled oligonucleotide specific for 5S rRNA. Below, the graph for the half-life determination is depicted.

As *sr7* is transcribed from the complementary strand to the essential *rpsD* gene, the entire *sr7* gene cannot be simply deleted. Therefore, we added the heterologous BsrF terminator [39] to the 3’ end of the *rpsD* gene and simultaneously replaced the *sr7* gene including its promoter p_*sr7*_ by the chloramphenicol resistance gene. The resulting strain DB104(*Δsr7*) was grown as above and induced with 0.5 M NaCl. Northern blotting confirmed that the knockout strain does not express the 185 nt SR7 (not shown). Therefore, this strain was employed below to analyse the function of SR7P.

### The small protein SR7P is synthesized in *B. subtilis*, and its amount increases under NaCl, ethanol and heat stress

In preliminary experiments, we had shown with a translational *lacZ* fusion under control of the constitutive strong promoter pIII [35] that the SD sequence of *sr7p* is functional in *B. subtilis* (results of β-galactosidase measurements are summarized in Fig. S2). To determine under which conditions the 39 aa protein SR7P is synthesized in *Bacillus subtilis*, strain DBSR7PF encoding a C-terminally FLAG-tagged peptide (SR7P_C-FLAG_) in its native locus was constructed as described in *Materials and Methods*. The SR7 transcriptional terminator was replaced by the heterologous BsrF terminator [39]. To confirm the synthesis of SR7P_C-FLAG_, 100 ml of strain DBSR7PF were grown in complex TY medium, not induced or subjected to different stresses for 15 min (NaCl, ethanol and acid stress, heat-shock) and crude extracts prepared by sonication. After equilibration to the same protein amounts, they were passed through M2 anti-FLAG columns, elution fractions pooled, precipitated, separated in a 15 % SDS-PAA gel and subjected to Western blotting using M2-anti FLAG antibodies. In non-stressed DB104, a weak protein band could be visualized at ≈ 6.5 kDa (Fig. 3A) corroborating synthesis of SR7P_C-FLAG_ in *B. subtilis*. Without the column step, a protein of nearly the same size that unspecifically interacted with the FLAG antibodies masked the SR7P signal. Threefold higher amounts of SR7P_C-FLAG_ were detected under ethanol stress, and twofold higher amounts under salt stress and heat shock (Fig. 3A). This increase was lower than expected, but most probably due to synthesis of certain amounts of SR7P already under non-stress conditions from the processing product of the SigA-dependent RNA species. The SR7P signal obtained after acid stress was very weak, but this seems to be an experimental artefact as we observed decreased amounts of all proteins after acid stress (not shown).

**Fig. 3.**
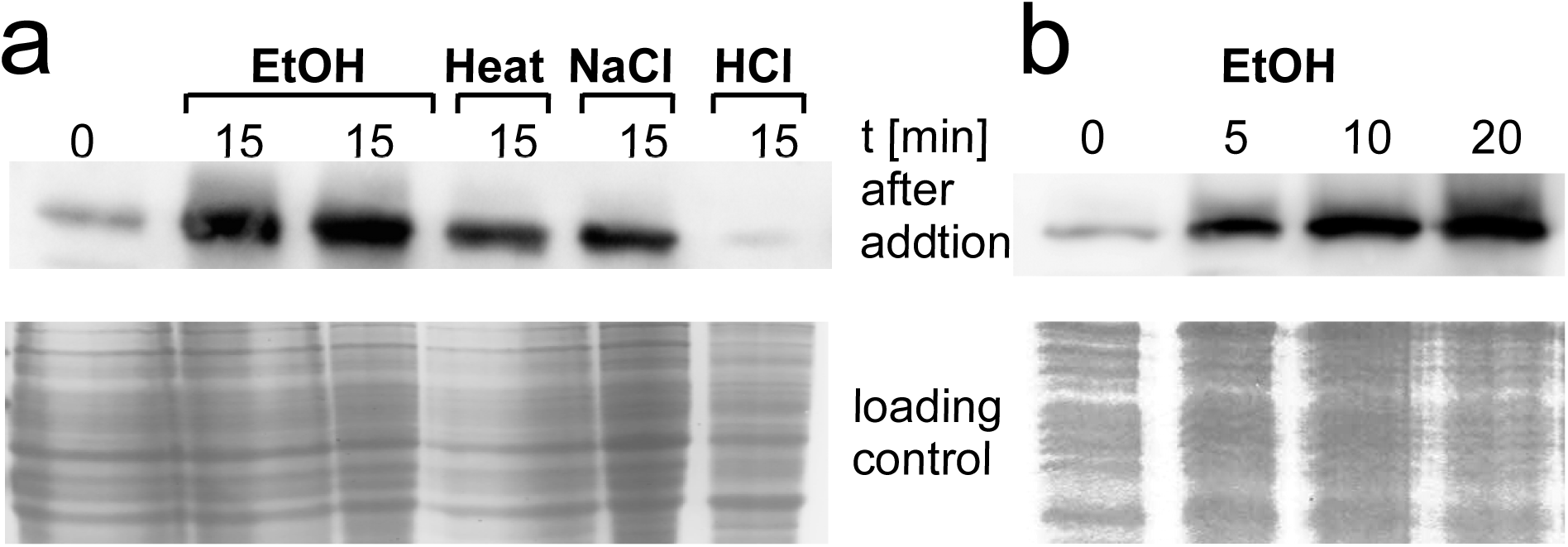
SR7P is synthesized in *B. subtilis* under non-stress and stress conditions. (a) *B. subtilis* strain DBSR7PF was grown in TY, treated for 15 min with 0.5 M NaCl, 4 % ethanol or 48 °C (heat shock), subjected to sonication, crude extracts equilibrated to the same total protein amounts and passed through anti-FLAG M2 columns and concentrated as described in *Materials and Methods*. Aliquots were loaded onto 15 % SDS PAA gels and subjected to Western blotting with anti-FLAG antibodies as described in *Materials and Methods*. Equilibrated crude extracts were used as loading control. (a) Comparison of the amounts of SR7P under non-stress and stress conditions. (b) Increase of SR7P amounts over time during ethanol stress.

In an additional experiment, cultures were subjected to ethanol stress, time samples taken after 5, 10 and 20 min and analysed as above. Already 10 min after ethanol addition, the highest amount of SR7P was observed (Fig. 3B). This is in good correlation with a SigB-dependent response.

### SR7P *interacts with enolase*

To identify potential interaction partners of SR7P, coelution experiments were performed. To this end, strain DBSR7PF expressing SR7P_C-FLAG_ was grown in 0.8 l complex TY medium. Strain DB104 expressing the untagged SR7P served as negative control. Crude extracts from both cultures were prepared (see *Materials and Methods*), applied to M2 anti-FLAG columns and elution fractions analysed on 15 % SDS PAA gels. After staining, one band of about 50 kDa was visible in elution fractions two and three from DBSR7PF, whereas no bands were detected in the negative control (Fig. 4A). The 50 kDa band was excised, subjected to tryptic digestion followed by mass spectrometry analysis and identified to be *B. subtilis* enolase (Eno). To confirm the SR7P_C-FLAG_ -Eno interaction, a reciprocal experiment was performed using strain DBSR7PFE encoding both N-terminally Strep-tagged enolase (Eno_N-Strep_) and in addition SR7P_C-FLAG_ from their native loci. Cultures were grown under the same conditions as above, crude extracts prepared, applied to Strep-Tactin columns, eluted with desthiobiotin, separated on 15 % SDS-PAA gels and stained (Fig. 4B). Subsequent Western blotting with M2 anti-FLAG antibodies revealed a direct correlation between the amount of eluted Eno_N-Strep_ and the amount of co-eluted SR7P_C-FLAG_ (Fig. 4B).

**Fig. 4.**
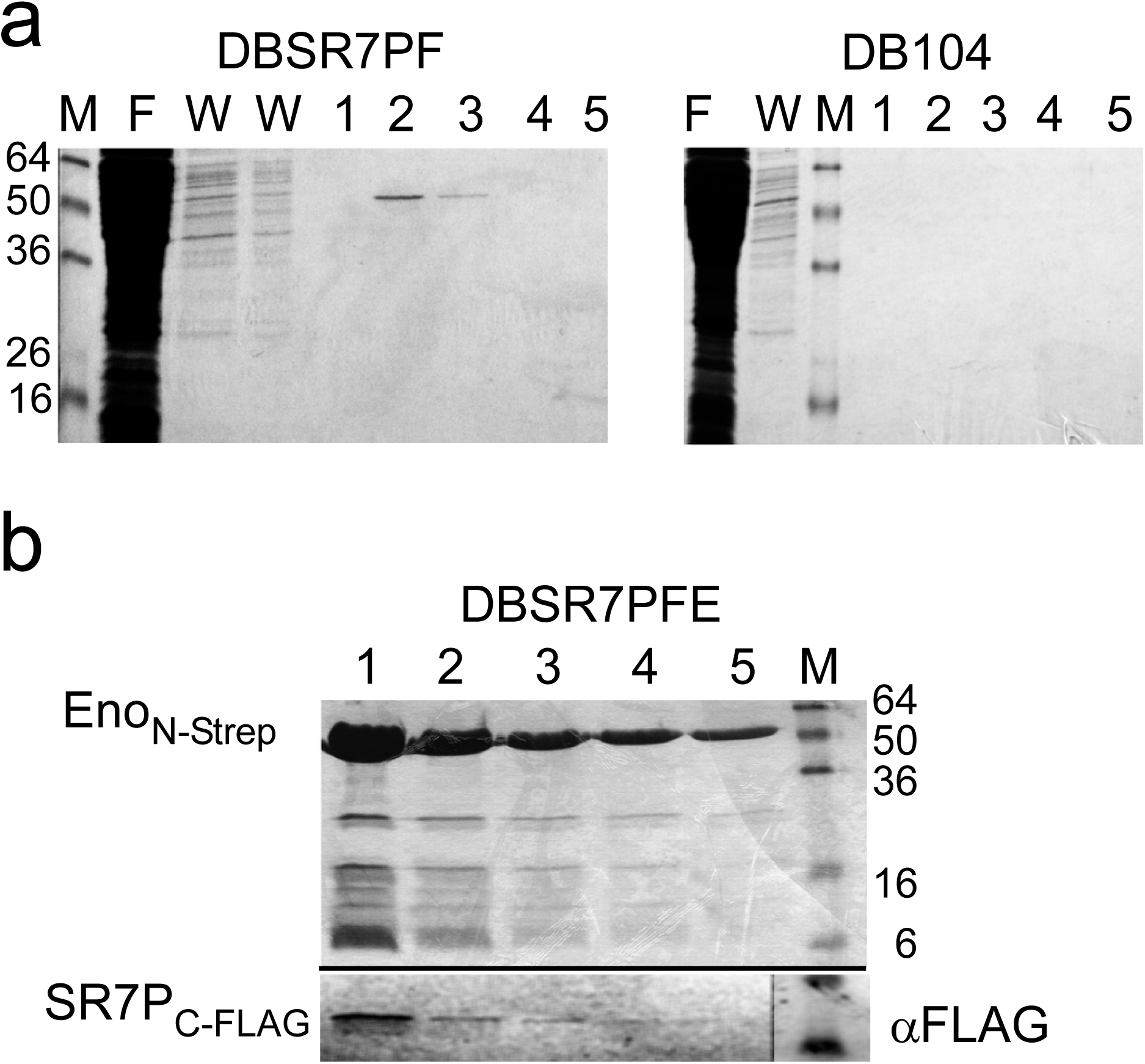
Enolase co-purifies with SR7P. *B. subtilis* strains DBSR7PF and DB104 were grown in 800 ml TY, crude protein extracts prepared as described in *Materials and Methods* and passed through anti-FLAG M2 columns. 25 μl of each 500 μl elution fraction were separated on 15 % SDS-PAA gels. (a) Coomassie-stained gels with the elution fractions from DBSR7PF and DB104. The visible ≈50 kDa band in elution fractions 2 and 3 of DBSR7PF gel was cut out, subjected to tryptic digestion and identified by mass-spectrometry to be enolase. (b) Reverse experiment: *B. subtilis* strain DBSR7PFE expressing Eno_N-Strep_ and SR7P_C-FLAG_ from the chromosome was grown as in (a), crude extracts prepared and passed through a Strep-Tactin column. 25 μl of each 500 μl elution fraction were separated as in (a) and subjected to Western blotting with α-FLAG antibodies to detect co-elution of SR7P_C-FLAG_.

To rule out that enolase might interact with any FLAG-tagged protein, strain DB104 expressing another FLAG-tagged small protein, SP2184_C-FLAG_, was employed in a co-elution experiment with Eno_N-Strep_ as in the previous experiment. Elution fractions from the Strep-Tactin-column displayed Eno_N-Strep_ on the stained gel, whereas no SP2184_C-FLAG_ was detected in the Western blot (Fig. S3). As a positive control for Western blotting and the functionality of the antibody served DBSR7PF expressing *sr7p*_*C-FLAG*_. To exclude a hypothetical interaction of the FLAG-tag with the Strep-Tag or the Strep-Tactin column, an additional control experiment was performed: The *gapA*_*C-Strep*_ gene [33] was integrated into strain DBSR7PF expressing *sr7p*_*C-FLAG*_. After preparing crude extracts as above and separating elution fractions from the Strep-Tactin column in SDS-PAA gels, GapA_C-Strep_ could be visualized on the stained gel, but no FLAG-tagged protein was detectable on the Western blot (Fig. S3).

Taken together, these experiments demonstrate that SR7P interacts with *B. subtilis* enolase.

### SR7P_C-FLAG_ co-elutes RNase Y bound to enolase but not phosphofructokinase PfkA

Recently, we have discovered that *B. subtilis* GapA binds RNase J1 and RNase Y [24,33]. By Western blotting we detected in GapA preparations from *B. subtilis* DB104 small amounts of RNase Y (one molecule RNase Y in 100 molecules GapA). Far Western blotting confirmed that RNase Y can bind GapA and vice versa. As enolase has been reported to interact with both RNase Y and phosphofructokinase PfkA in the *B. subtilis* degradosome [36,40] we wanted to find out whether SR7P_C-FLAG_ coelutes RNase Y or PfkA bound to enolase. To enable detection of PfkA in Western blots, strain DBPF_C-His_ encoding PfkA_C-His6_ and SR7P_C-FLAG_ in their native loci was constructed. This strain was used for a co-elution experiment as above, followed by Western blotting with antibodies against native RNase Y and against the His-Tag. As shown in Fig. 5, enolase was co-eluted as before (Fig. 5A) and RNase Y (58 kDa) could be identified in the same elution fractions (Fig. 5B). By contrast, no co-eluted phosphofructokinase could be detected (Fig. 5C). Therefore, we conclude that SR7P_C-FLAG_ coelutes enolase carrying RNase Y but not phosphofructokinase.

**Fig. 5.**
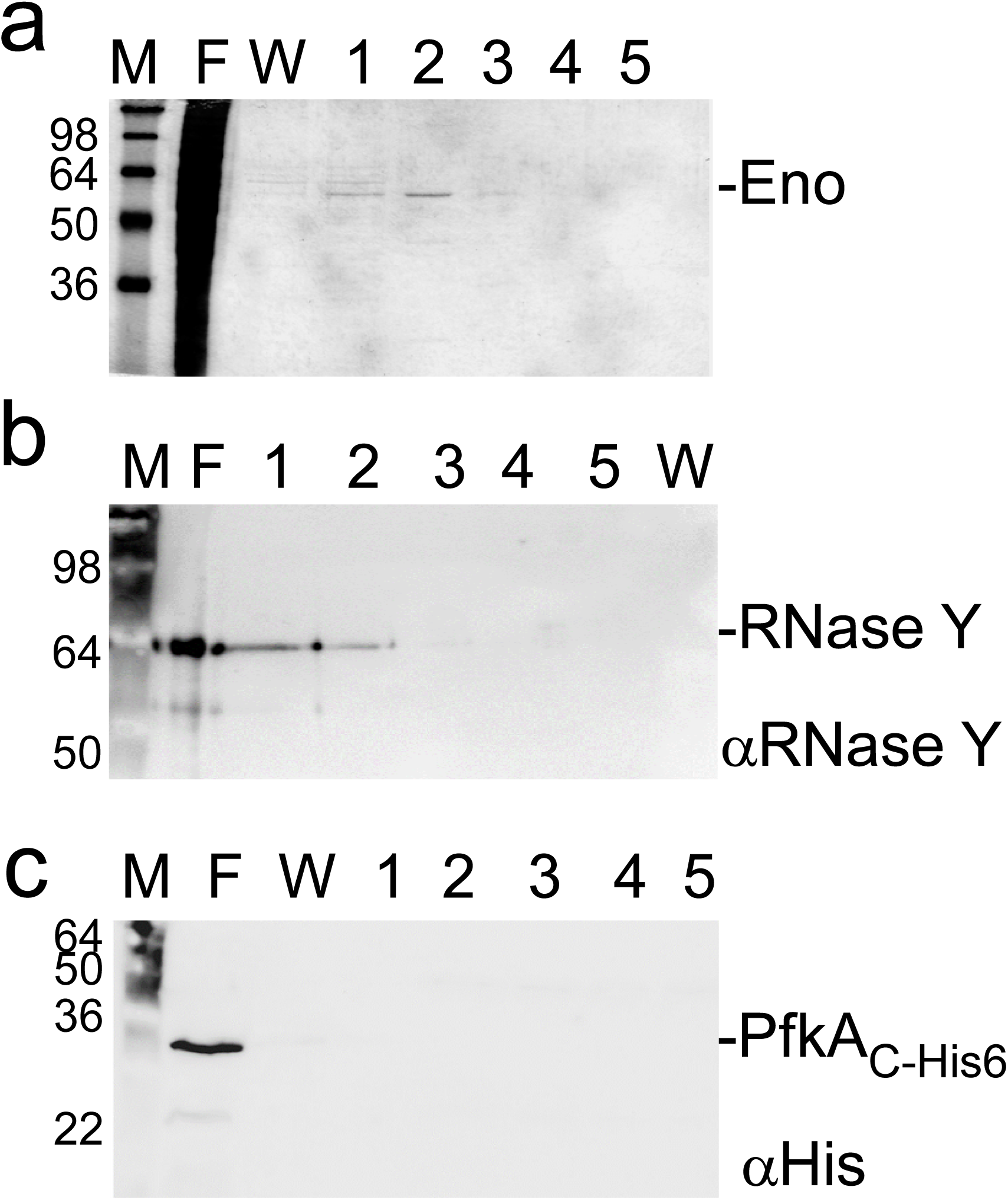
SR7P co-elutes RNase Y bound to enolase, but not phosphofructokinase PfkA. *B. subtilis* DBPF_C-His_ expressing both His-tagged PfkA and SR7PC_-FLAG_ from their native loci was grown in TY medium until stationary phase, crude protein extracts prepared as described in *Materials and Methods* and separated through FLAG M2-columns. Elution fractions containing co-eluted enolase were run on 15 % SDS-PAA gels (a) and subjected to Western blotting with anti-RNase Y antibodies (b) or anti-His-tag antibodies (c).

### Far Western blotting confirms the interaction between RNase Y or SR7P with enolase but excludes a direct SR7P-RNase Y interaction

To confirm the SR7P-enolase interaction by an independent method and to exclude a direct SR7P-RNase Y interaction, Far Western blotting was employed. First, we used Eno_N-Strep_ isolated from the wild-type and from the *Δsr7* strain as targets and RNase Y_C-His6_ as bait (Fig. 6A). Detection was performed with antibodies against native RNase Y, which proved to be specific (control, central panel). RNase Y bound specifically both Eno preparations (right panel), indicating that the Strep-tag does not interfere with binding. In the second Far Western blot (Fig. 6B), we used RNase Y_C-His6_ as target and Eno_N-Strep_ with or without SR7P as bait, and detection was with anti-Strep-tag antibodies, which were also specific (control panel). As expected, Eno_N-Strep_ and Eno_N-Strep_ /SR7P bound to RNase Y (right). To exclude a direct interaction between SR7P and RNase Y we run RNase Y_C-His6_ and – as positive control – Eno_N-Strep_ on a gel and used SR7P_C-FLAG_ purified from IPTG-induced *E. coli* strain BL21DE3(pDRSR7P), which does not contain RNase Y, as bait (Fig. 6C). Whereas a strong signal for SR7P_C-FLAG_ bound to enolase was visible, no signal for an interaction with RNase Y was detectable.

**Fig. 6.**
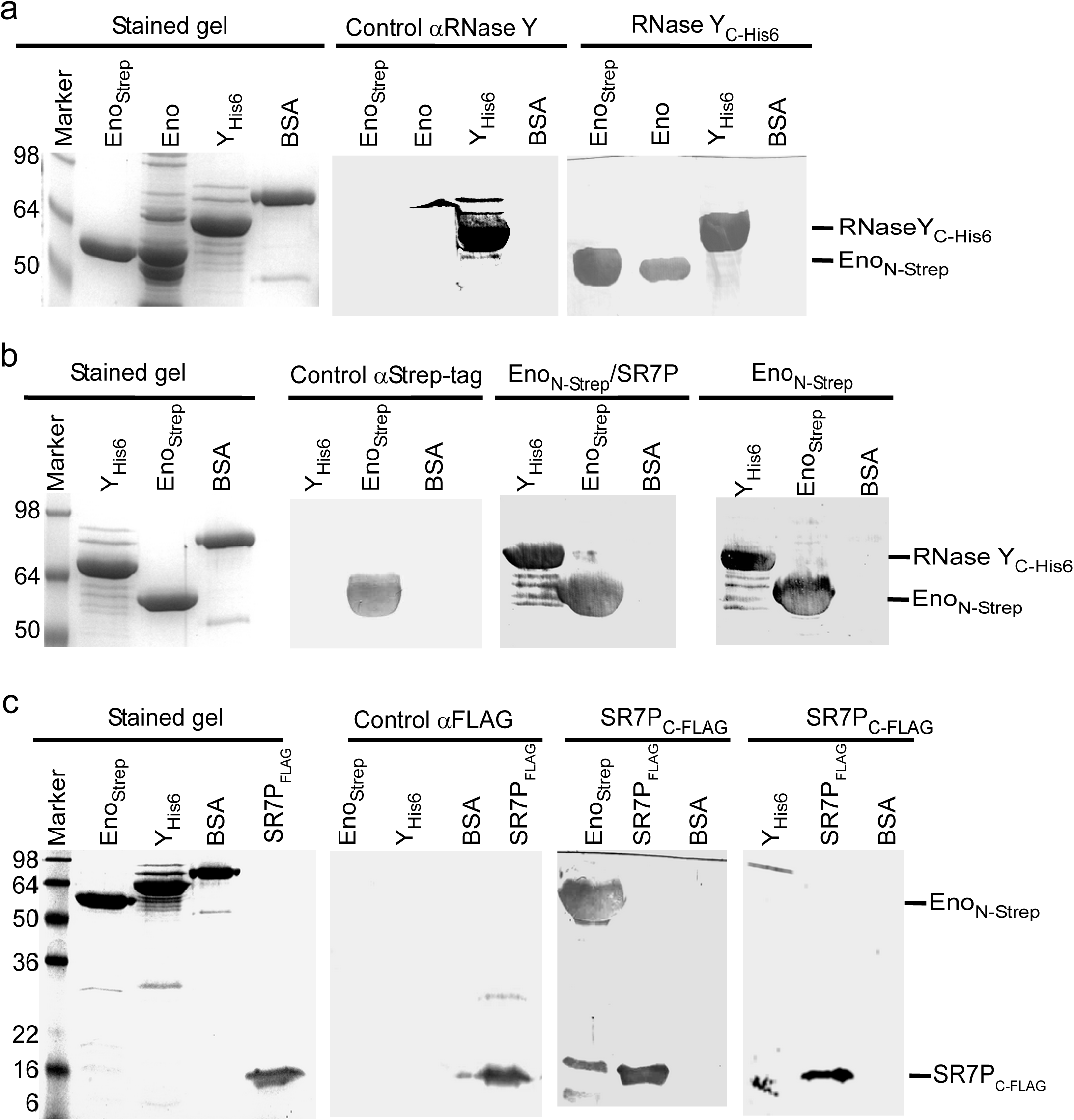
Far Western blotting. Representative blots of two independently performed experiments are shown. Proteins were separated on 10 % or 15 % SDS-PAA gels and either stained with Coomassie or blotted on PVDF membrane as described in *Materials and Methods*. RNase Y_C-His6_ was purified from *E. coli*, Eno_N-Strep_ was purified from *B. subtilis* and untagged Eno was co-purified with SR7P_C-FLAG_ from *B. subtilis*. (a) After blocking all blots were incubated with PBST gelatine (control) or 10 ml PBST-gelatine containing 170 µg RNase Y_C-His6_. RNase Y binding was detected with native antibodies against RNase Y. RNase Y was able to bind Eno and Eno_N-Strep_. (b) Far Western Blot as in (a) except that blots were incubated with 100 µg Eno_N-Strep_ purified from either *B. subtilis* GP1215 or *B. subtilis* DBSR7PFE. Eno_N-Strep_ binding was detected with mouse anti-Strep-tag antibodies. (c) SR7P_C-FLAG_ purified from *E. coli* strain BL21DE3(pDRSR7P) was used to confirm the SR7P-enolase interaction and to exclude a direct SR7P-RNase Y interaction. SR7P_C-FLAG_ binding was detected with anti-FLAG antibodies.

Taken together, Far Western blotting corroborated the enolase-SR7P and the enolase-RNase Y interactions discovered in co-elution experiments and ruled out a direct SR7P-RNase Y interaction.

### Enolase co-eluted with SR7P_C-FLAG_ carries significant amounts of RNase Y

To investigate if SR7P influences the binding of RNase Y to enolase, we performed co-elution experiments with two strains both encoding Eno_N-Strep_ at its native locus. One strain contained in addition the *sr7p*_C-FLAG_ gene, in the other strain the entire *sr7* gene was replaced by the *cm*^*R*^ gene. Eno_N-Strep_ co-eluted by SR7P_C-FLAG_ was compared to Eno_N-Strep_ eluted from the *Δsr7* strain through a Strep-Tactin column. The elution fractions were subjected to Western blotting. As shown in Fig. 7A, in the presence of SR7P, enolase carried about 10 times more RNase Y than in its absence. In fact, an approximately 1 : 1 ratio of enolase and RNase Y was detected in the presence of SR7P. These data indicate that SR7P increases the amount of RNase Y bound by enolase or promotes the enolase-RNase Y interaction.

**Fig. 7.**
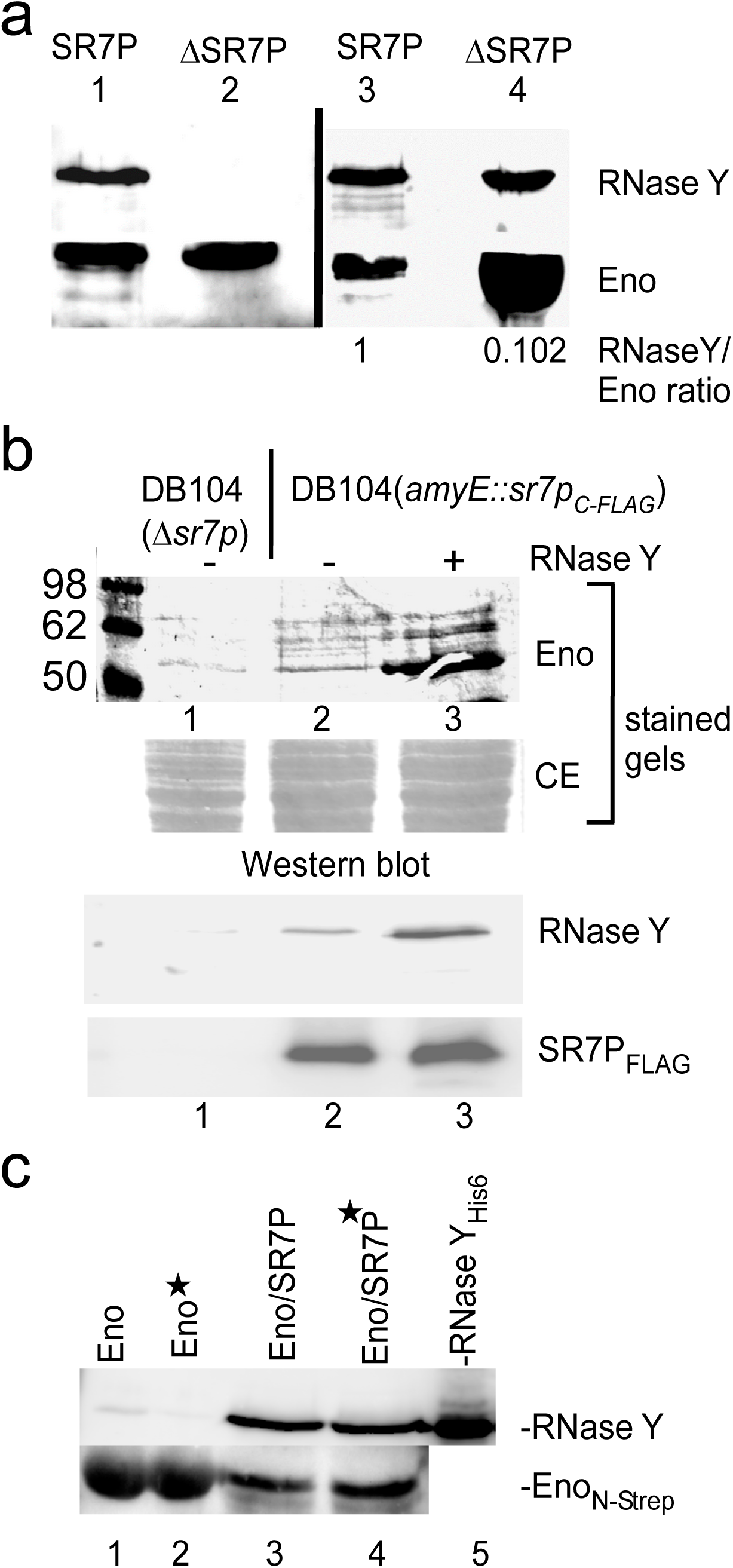
Enolase-SR7P co-elutes significant amounts of RNase Y and RNase Y promotes binding of enolase to SR7P. (a) Co-elution experiment performed as in Fig. 4 with strains DBSR7PFE and DB104(*Δsr7*, eno_N-Strep_). Elution fractions were subjected to Western blotting with anti Strep-tag antibodies (for enolase) and mouse antibodies against native RNase Y employing chemiluminescence. Two parallels are shown. In lanes 1 and 2, identical amounts of enolase were loaded (based on a Coomassie-stained gel). Here, only in the presence of SR7P, an RNase Y signal is detectable. In lanes 3 and 4, a 9-fold higher amount of enolase had to be loaded from the *Δsr7* strain to yield an RNase Y signal of the same intensity. (b) Coelution experiment with DB104(*Δsr7*) and DB104 expressing IPTG-inducible SR7P_C-FLAG_ from a single copy in the *amyE* locus. Cells were grown in TY, induced for 30 min with IPTG, crude extracts prepared, equilibrated to the same protein concentration and passed through anti-FLAG M2 columns. +, 100 μg RNase Y_C-His6_ purified from *E. coli* were added to the crude extract before column loading. Elution fractions were separated on a 15 % SDS-PAA gel and subjected to Western blotting with anti-RNase Y or anti-FLAG-tag antibodies. (c) Role of RNA in the enolase-RNase Y interaction. Co-elution experiments were performed as described in Fig. 4, but with DB104 (*eno*_*N-Strep*_, *sr7p*_*His10*_) and 50 % of the crude extracts were treated for 10 min with 10 mg/ml RNase A at room temperature (*) prior to loading onto Ni-agarose columns. Elution fractions were subjected to Western blotting with antibodies against native RNase Y and the Strep-tag, respectively. Lanes 1 and 2, Eno_N-Strep_ purified from DB104(*eno*_*N-Strep*_, *Δsr7*); lanes 3 and 4, Eno_N-Strep_ purified from DB104 (*eno*_*N-Strep*_, *sr7p*_*His10*_); lane 5, 0.5 pmol RNase Y_C-His6_ purified from *E. coli*.

### RNase Y promotes the interaction between enolase and SR7P

When we added RNase Y_C-His6_ purified from *E. coli* to a crude-protein extract obtained from a *B. subtilis* strain expressing an IPTG-inducible SR7P_C-FLAG_ before loading it onto an anti-FLAG M2 column, we observed in the elution fractions a significant increase in the amount of enolase bound to SR7P_C-FLAG_ (Fig. 7B, lane 3 vs. 2) although the amount of SR7_C-FLAG_ detected in Western blots was identical. The background bands in the stained gels above the enolase band are due to treatment with IPTG. Since we have shown in Far-Western blots (Fig. 6C) that SR7P cannot directly bind RNase Y, i.e. does not bridge the enolase-RNase Y interaction, this result suggests that RNase Y facilitates the binding of enolase to SR7P. The results displayed in Figs. 7A and B are indicative of an “induced-fit” mechanism.

### RNA does not bridge the interaction between SR7P, enolase and RNase Y

To investigate if RNA is required for the SR7P/enolase and enolase/RNase Y interactions, a co-elution experiment was conducted using two *B. subtilis* strains, both expressing *eno*_*N-Strep*_ from the chromosome and one lacking the *sr7* gene, the other expressing *sr7p*_*C-His10*_. Crude protein extracts – 50 % treated with RNase A and 50 % untreated – were passed through Ni-NTA agarose columns. Afterwards, elution fractions were analysed by Western blotting with anti-RNase Y antibodies and anti Strep-tag antibodies (Fig. 7C). As shown in lanes 3 and 4, identical amounts of enolase and RNase Y were co-eluted with SR7P_His10_ in RNase A-treated and in untreated extracts. This confirmed the Far Western blotting data that SR7P can bind enolase without the help of RNA (Fig. 6C). Furthermore, it also corroborated that the enolase/RNaseY/SR7P complex forms without the help of RNA as scaffold. However, when we purified Eno_N-Strep_ from a *Δsr7* strain through a Strep-Tactin column and assayed co-eluted RNase Y in Western blots (lanes 1 and 2), we observed about 20 % to 30 % less RNase Y in the RNase A-treated samples. This suggests that although enolase interacts with RNase Y in the absence of SR7P (Fig. 6A and B, and [40]) the interaction is to a minor degree supported by RNA.

### In vitro *degradation of RNase Y substrates* yitJ *5’ UTR and* rpsO *mRNA*

A number of RNAs have been reported to be substrates of RNase Y [41]. Among them is the 5’ UTR of the riboswitch *yitJ*, which had been confirmed both *in vivo* and *in vitro* to be an RNase Y substrate [42]. All other substrates have been characterized *in vivo* in Northern blots, but none of them has been analysed *in vitro* with purified RNase Y. Therefore, we set up an *in vitro* degradation assay with 5’ ^32^P-labelled substrate RNA, RNase Y_C-His6_ purified from *E. coli* and Eno_N-Strep_ purified from *B. subtilis sr7p*_C-FLAG_ or *Δsr7* strains. Eno_N-Strep_ contains the co-eluted RNase Y. When we used enolase carrying SR7P_C-FLAG_, degradation of *yitJ* RNA was visible with 0.3 pmol Eno_N-Strep_/RNase Y/SR7P. By contrast, even 12 pmol of Eno_N-Strep_/RNase Y purified from the *Δsr7* strain were not sufficient for a significant increase in *yitJ* RNA degradation (Fig. 8A). This result suggests that SR7P does not only enhance binding of RNase Y to enolase but, in addition, it increases the enzymatic activity of RNase Y in this triple complex. Previous Northern blots showed that *rpsO* mRNA is also a substrate of RNase Y [43], however, no *in vitro* cleavage had been studied previously and no transcription start site published. Since we observed in Northern blots three *rpsO* RNA species, the second of which was the main RNase Y target (see below), we determined their 5’ ends by primer extension (Fig. S4). For our *in vitro* degradation studies, we used the *in vitro*-synthesized 5’ labelled 386 nt *rpsO* mRNA as substrate (Fig. 8B). First, we employed different amounts of RNase Y_C-His_ purified from *E. coli* to analyse the degradation pattern of *rpsO* mRNA (left). Then, we used enolase that does not contain RNase Y as well as SR7P purified from *E. coli* as negative controls to ensure that they do not have intrinsic RNase activities (left). When we compared the activity of RNase Y in the Eno_N-Strep_ /RNase Y/SR7P complex with that in the Eno_N-Strep_/RNase Y complex purified from the *Δsr7* strain, we found it to be significantly more active. Like in the case of *yitJ* RNA, RNase Y bound to 0.3 pmol Eno_N-Strep_ in the triple complex was sufficient for complete disappearance of the full-length *rpsO* species whereas no substantial degradation was observed with even 3.2 pmol SR7P-free Eno_N-Strep_ /RNase Y. Interestingly, when SR7P purified from *E. coli* (SR7P_E_) was added afterwards to 0.8, 1.6 or 3.2 pmol Eno_N-Strep_ /RNase Y, at 3.2 pmol, full-length *rpsO* RNA was degraded almost completely (last lane), and the same distinct pattern of degradation products was observed as with Eno_N-Strep_/RNase Y/SR7P at 0.1 pmol enolase. This indicates that even the subsequent addition of purified SR7P can improve degradation of *rpsO* RNA by enolase-bound RNase Y, although the complex formed *in vivo* seems to be more active. Surprisingly, 1 pmol of RNase Y purified from *E. coli* were less active in degrading *rpsO* mRNA than the small amount of RNase Y present in the complex with 0.1 pmol enolase and SR7P. This again suggests that SR7P might increase the enzymatic activity of RNase Y bound to enolase.

**Fig. 8.**
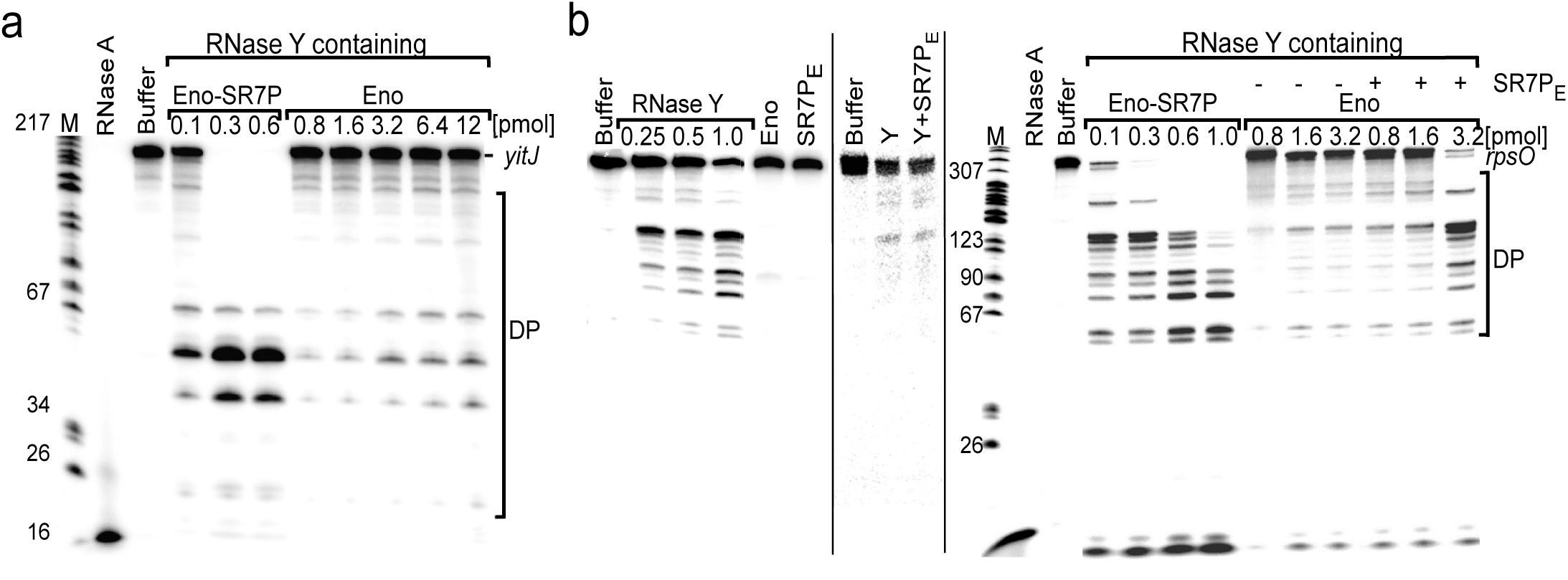
SR7P affects RNA degradation *in vitro*. RNA degradation assays with 5’-labelled 5’ UTR of *yitJ* RNA and *rpsO* mRNA (386 nt) were performed as described in *Materials and Methods*. Protein-free buffer was used as negative control and 10 pmol RNase A as positive control. 5’-labelled pBR322xMspI served as size marker. The protein amounts (pmol enolase) used, full-length substrate RNA and degradation products (DP) are indicated. (a) Degradation of *yitJ* 5’ UTR (b) Degradation of *rpsO* mRNA. Left and middle: separate gels with controls: Degradation of *rpsO* mRNA with RNase Y (pmol are indicated) purified from *E. coli* shows the identical degradation pattern as on the right side. Degradation with enolase (Eno_P_) purified via Strep-Tactin columns followed by ion exchange chromatography and gel filtration to remove bound RNase Y, or with SR7P_E_ purified from *E. coli* show that neither RNase Y-free enolase nor SR7P are able to cleave *rpsO* mRNA. The addition of SR7P_E_ does not affect the cleavage by RNase Y. Right: cleavage of *rpsO* mRNA by RNase Y copurified with enolase from a *B. subtilis* strain expressing SR7P_C-FLAG_ (lanes 4-7) and from a *Δsr7* strain (lanes 8-13). Lanes 11-13, subsequent addition of 3 pmol SR7P_E_ increased the activity of 3.2 pmol enolase-bound RNase Y. Eno, Eno_N-Strep_ purified from *B. subtilis*; Eno-SR7P, Eno_N-Strep_ with SR7P_C-FLAG_ co-purified from *B. subtilis*.

From these *in vitro* data we reason that SR7P promotes degradation of *yitJ* and *rpsO* RNAs by enolase-bound RNase Y.

### *Analysis of effects of SR7P, enolase and RNase Y on* rpsO *mRNA* in vivo

To analyse the effects of SR7P, enolase and RNase Y on the half-life of *rpsO* mRNA *in vivo*, we performed Northern blotting experiments with wild-type DB104 and isogenic *Δrny, Δeno* and *ΔylbF* strains grown in TY until stationary phase. The *ylbF* knockout strain was included, because recently the Y-complex composed of YlbF, YmcA and YaaT has been discovered to affect the degradation of many mRNAs by RNase Y [44]. As expected for an RNase Y substrate, the half-life of *rpsO* mRNA was with ≈29 min at least tenfold longer in the absence of RNase Y (Fig. 9A). In the absence of enolase, the half-life increased about 2.5-fold, and in the absence of the Y complex component YlbF about threefold. As we observed twofold higher amounts of SR7P after salt stress (Fig. 3B) we determined the half-life of *rpsO* mRNA in the wild-type and the isogenic *Δsr7* strain after treatment with 0.5 M NaCl. We did not employ ethanol stress, because ethanol had also an effect on the abundance of some RNase Y targets *in vivo* [45,46]. With 2.32 vs. 2.93 min we observed a slight, but reproducibly higher half-life in the absence of SR7P (Fig. 9B). To corroborate this effect, we constructed strain DB104 (*amyE::sr7p*_*C-FLAG*_ spec^R^) for inducible overexpression of *sr7* from the chromosomal *amyE* locus. When we induced this strain with IPTG, the *rpsO* RNA half-life decreased from 4.05 (DB104) to 3.12 min (Fig. 9C). This was again a slight effect, which was, however, reproducible with four biological replicates.

**Fig. 9.**
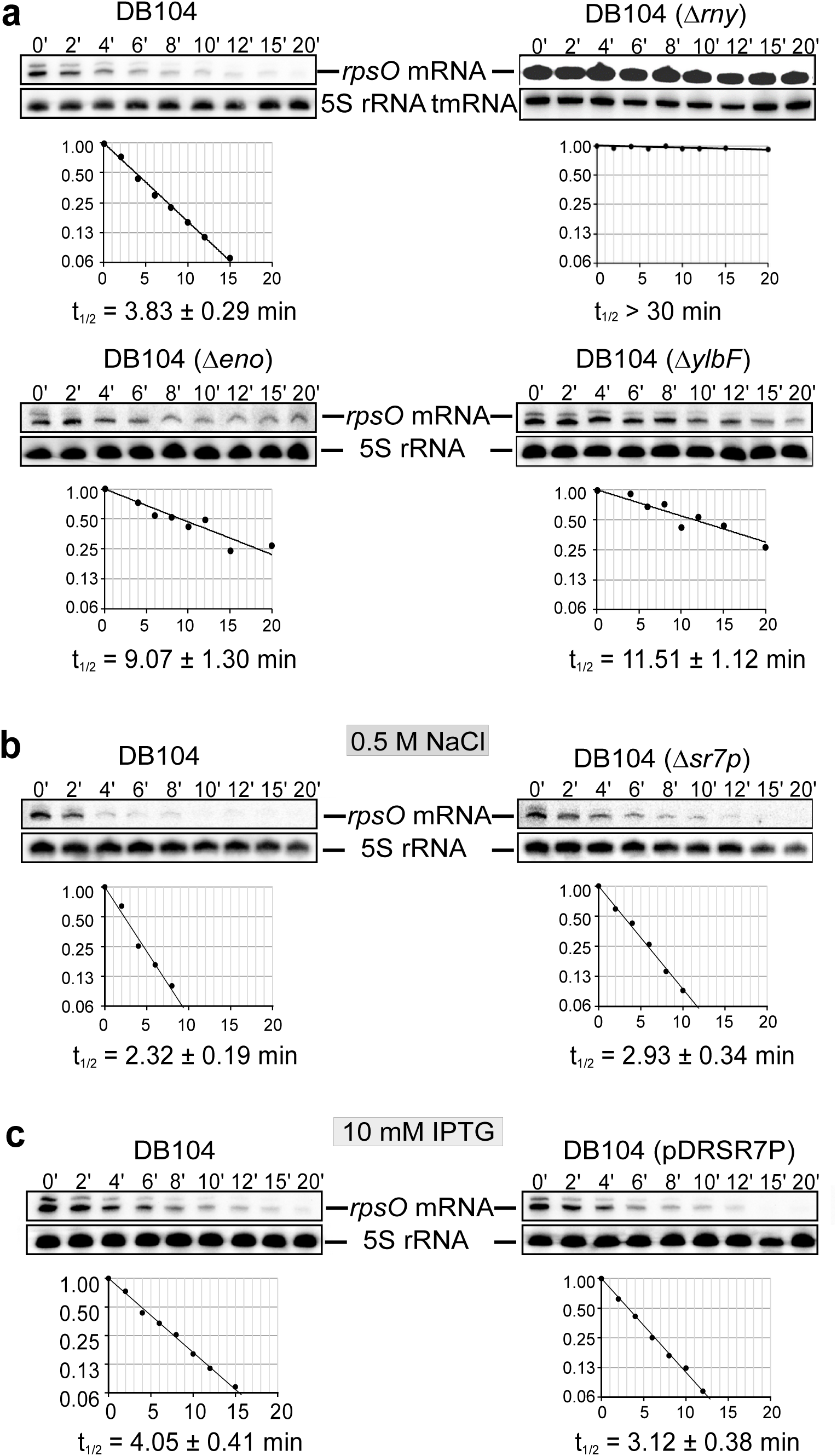
Effect of SR7P, RNase Y, enolase and YlbF on *rpsO* mRNA stability in *B. subtilis*. *B. subtilis* strains were grown in TY medium, time samples taken as described in *Materials and Methods* and used for the preparation of total RNA. RNA was separated on 6 % denaturing PAA gels, blotted onto nylon membrane. Detection of *rpsO* mRNA and reprobing were performed as in Fig. 2. Autoradiograms of the Northern blots are shown. (a) The half-life of *rpsO* mRNA was determined in DB104 and the isogenic *Δrny, Δeno and ΔylbF* strains after rifampicin addition. (b)The half-life of *rpsO* mRNA was determined in DB104 and DB104(*Δsr7*) after 30 min treatment with 0.5 M NaCl followed by rifampicin addition. (c) The half-life of *rpsO* mRNA was determined in wild-type strain DB104 and in DB104(*amyE::sr7p*_*C-FLAG*_ spec^R^) (inducible overexpression of *sr7*) 30 min after addition of 10 mM IPTG followed by rifampicin addition.

### Half-life of SR7 in the presence or absence of RNase Y, enolase and YlbF

As it is also not excluded that the stability of SR7 itself is dependent on RNase Y or the Y complex, we determined the stability of SR7 in the isogenic *Δrny* and *ΔylbF* strains grown in TY until stationary phase. Fig. S1C shows that the stability of SR7 is neither affected by RNase Y nor by YlbF.

### SR7P impacts the survival of *B. subtilis* after ethanol, acid and heat stress

To uncouple the biological function of SR7 from that of the encoded sprotein SR7P, wild-type strain DB104 and the isogenic mutant strain DBSR7P-1 containing a start-to stop-codon mutation in the *sr7* gene but expressing SR7 under stress conditions were grown in complex TY medium until stationary phase and subjected to ethanol stress (12 % ethanol), acid stress (pH 4.0), heat stress (57 °C), salt stress (4 M NaCl) and manganese stress (15 mM Mn^2+^) as described in *Materials and Methods*. When cells were plated 30 min after stress induction, the survival of the wild-type strain was ≈fourfold that of the mutant strain after ethanol stress, twofold after heat stress and 1.5- to twofold after acid stress (Fig. 10). By contrast, the presence of SR7P did not affect survival after salt or manganese stress significantly.

**Fig. 10.**
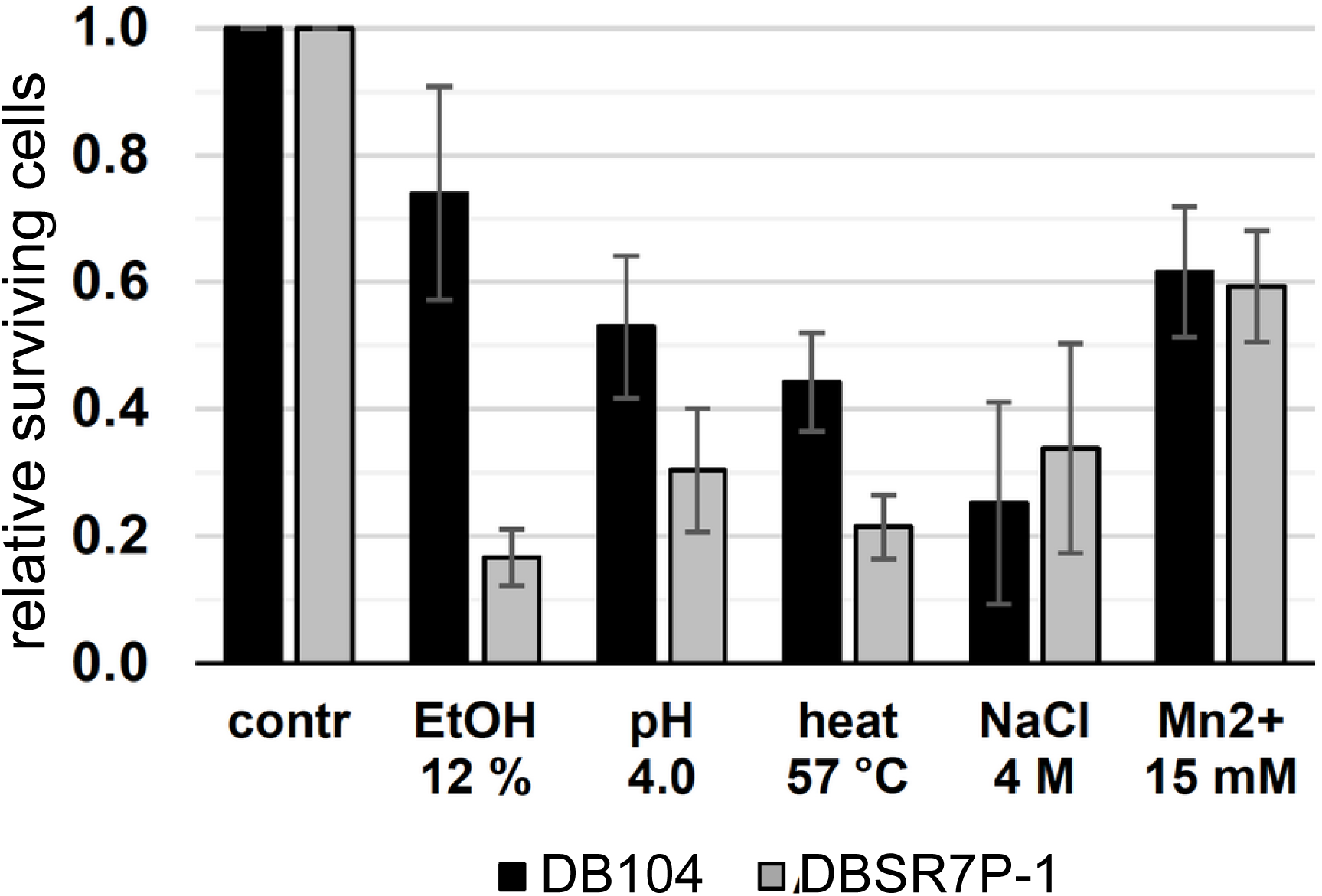
Impact of SR7P on survival after stress. Survival experiments were performed as described in *Materials and Methods* with wild-type strain DB104 and the isogenic DBSR7P-1 strain that contains a start- to stop codon mutation in the *sr7* gene. The results of 7 independent experiments with standard deviations are shown.

### SR7P is highly conserved among 10 Bacillus species

To ascertain if the 39 aa SR7P is restricted to *Bacillus subtilis*, we searched for sproteins with the same or similar aa sequences in other annotated genomes. SR7P homologues were only found in *B. subtilis* strains and in 9 other species belonging to the genus Bacillus. Neither in the genomes of other firmicutes nor in those of other Gram-positive or Gram-negative bacteria, a small protein with a similar primary sequence was found to be encoded. Fig. 11 presents an alignment of the aa sequences of these SR7P homologues. In all cases, an almost identical stretch of 20 aa is present in the N-terminal half of the protein, whereas the C-terminal half reveals more differences. In contrast to *B. subtilis* SR7P, all other homologues contain three additional hydrophilic aa near their C-termini and are, therefore, 42 aa long. When we assume that the SR7P-enolase interaction is conserved, the interaction surface is located most likely within the 20 conserved aa of SR7P.

**Fig. 11.**
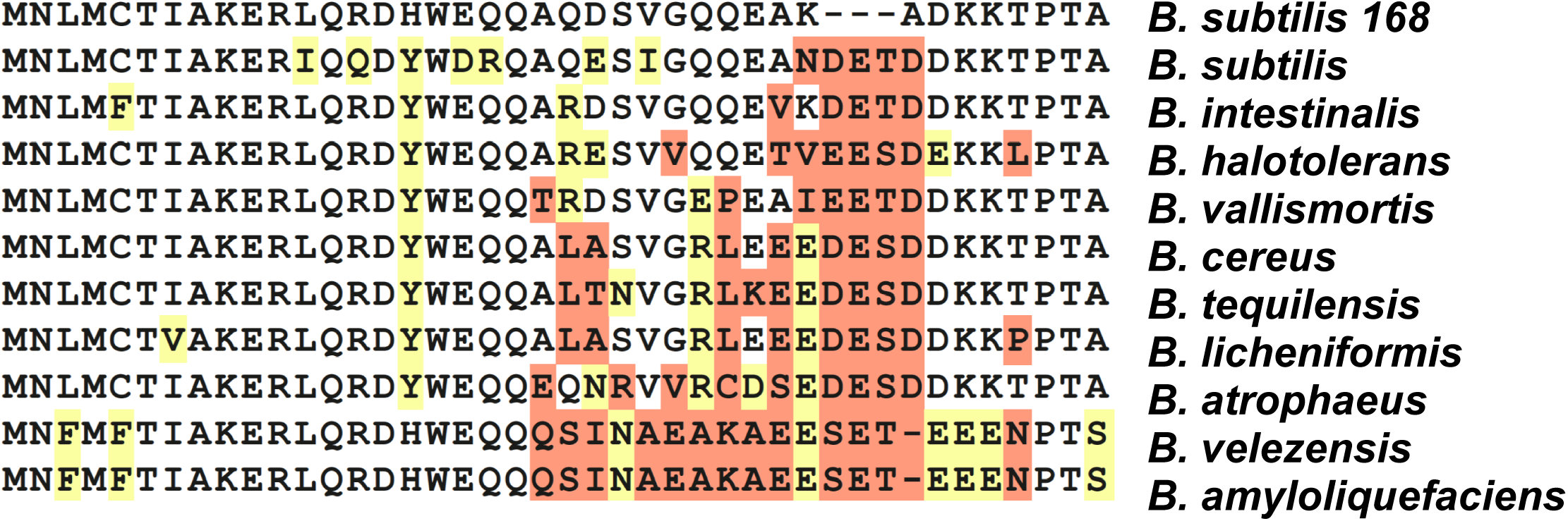
Alignment of SR7P homologues. Clustal Omega alignment of SR7P homologues. The SR7P sequences from 10 species were aligned. Only differences are highlighted; yellow, similar aa; orange, aa that differ either by charge or by hydrophobicity.

The peptide is predicted to form an α-helix. However, NMR data obtained so far with a purified SR7P (designated SP-12 in [47]) indicated that only 20.5 % of free SR7P is structured and 79.4 % is disordered. Interestingly, in predictions with the FuzPred algorithm for the bound form, the percentage of structured form increased to 41 % compared to 59 % disordered. Both CD and NMR structural analysis indicated a rather molten globule for free SR7P [47].

At DNA level, all homologous *sr7* genes are preceded by SigB-dependent promoters, and upstream of all peptide sequences SD sequences are located (Fig. S5). Only in *B. amyloliquefaciens* and *B. velezensis*, the distance between SD sequence and GTG start codon is with 14 nt rather long, but an alternative AG-rich sequence nearer to the start codon could also serve as RBS. In all cases, the start codon is GTG. In the 5’ half of the ORF, nt exchanges are found almost exclusively at wobble positions indicating that this is the most highly conserved region. All *sr7* genes carry Rho-independent transcription terminators at their 3’ ends. Consequently, the SigB-dependent SR7 homologues are also conserved at DNA level.

Only in *B. halotolerans, B. vallismortis* and *B. cereus*, the SR7P encoding genes are located downstream of the *tyrS* gene and convergently transcribed to the *rpsD* gene as in *B. subtilis* (see Fig. 1). However, not the entire genomes are annotated for the other species so far. Therefore, we cannot exclude a conserved gene arrangement.

## Discussion

The rapid degradation of mRNA is a vital process, since it contributes to the ability of bacteria to adapt to changing environmental conditions. This decay involves a number of endo- and exoribonucleases organized in a multiprotein complex, the so-called degradosome. The proposed *B. subtilis* degradosome was reported to contain the main endoribonuclease RNase Y, the main 3’-5’-exoribonuclease PnpA, RNases J1 and J2, helicase CshA, and the glycolytic enzymes Eno and PfkA [36]. However, this degradosome seems to have a more dynamic structure. In contrast to the *E. coli* degradosome, it cannot be isolated in the absence of cross-linking reagents [48]. Furthermore, not only Eno and PfkA, but also a third glycolytic enzyme, glyceraldehyde-3P-dehydrogenase A (GapA), affects the composition of this dynamic degradosome [33]: GapA interacts with RNase J1, and this interaction is improved by a small protein, SR1P, which is expressed under gluconeogenic conditions when the metabolic function of GapA is not required. Furthermore, the enzymatic activity of RNase J1 is enhanced in the GapA/RNase J1/SR1P complex compared to that of the unbound enzyme.

Here, we report on the discovery and characterization of another small protein in *B. subtilis*, the 39 aa SR7P. It is encoded on two RNAs, a constitutively transcribed and subsequently processed one of 260 nt and on a stress-induced SigB-dependent 185 nt RNA renamed as SR7 (formerly S1136, [37]). SR7P interacts with the glycolytic enzyme enolase. This interaction promotes binding of endoribonuclease RNase Y to enolase and significantly increases degradation of two known RNase Y substrates, *yitJ* RNA and *rpsO* mRNA, *in vitro*. In addition, *rpsO* degradation is also affected *in vivo* about threefold by the presence of enolase and to a minor, but reproducibly measurable extent, by SR7P (Fig. 9). Under salt stress, the twofold amount of SR7P is synthesized and bound by enolase, resulting in an increase of the enolase fraction carrying RNase Y and in turn, a slight increase in degradation of RNase Y substrates (Fig. 9). These data support that the Eno/SR7P-RNase Y interaction has a biological function. The small effect of inducible overexpression or deletion of *sr7* might be due to the fact that the bulk of its interaction partner enolase is located in the cytosol to fulfil its role in glycolysis, and only a small percentage is present in the degradosome where it interacts with RNase Y and PfkA. Furthermore, the effect of enolase deletion is higher than that of *sr7* deletion, because enolase moonlights in RNA degradation under a variety of conditions and SR7P only modulates this activity under specific stress conditions.

Binding of SR7P to enolase and formation of the SR7P/enolase/RNase Y complex are independent of RNA. Therefore, we can rule out a bridging function of RNA in the trimeric complex (Fig. 7C). Unexpectedly, binding of enolase to RNase Y in an *sr7* knockout strain was to a low extent supported by RNA (Fig. 7C). This is in contrast to what we found for the GapA-RNase Y or the GapA-RNase J1 interactions which were independent of RNA [33]. Furthermore, we could exclude that SR7P binds directly to RNase Y (Fig. 6C). We hypothesize that binding of SR7P alters the conformation of enolase and increases its binding affinity to RNase Y. In turn, the addition of RNase Y promoted the enolase-SR7P interaction (Fig. 7B). Since only 20.5 % of free SR7P are structured, and predictions indicated an increase of structure to 41 % upon binding to a partner molecule [47] our data support an “induced fit”-mechanism for the generation of a stable trimeric SR7P-enolase-RNase Y complex in the degradosome. This is in agreement with a large number of proteins in eukaryotes, but also in prokaryotes, comprising intrinsically disordered regions for which folding upon binding is an important mode of molecular recognition (rev. in [49]).

Already in 2011 it was reported that citrate alters the activity of *E. coli* PNPase by directly binding to the enzyme [50]. When Newman *et al*. found that citrate also influences the activity of *B. subtilis* enolase, they argued that glycolytic enzymes may act as sensors of nutritional stress and coordinate this stress with the RNA degrading machinery [40]. This would entail a decrease of global mRNA turnover under energy-limiting conditions. Our previous data on SR1P/GapA supported this idea: SR1P is only synthesized when glucose is exhausted and might also act as a sensor to link RNA degradation to the nutritional state of the cell [33]. Similarly, SR7P might act as a sensor to connect RNA turnover to different stresses: *sr7* transcription was induced under five stress conditions, and a two- to threefold increase of SR7P under stress was found (Figs. S1 and 3). Under salt stress we observed a small, but reproducible effect of SR7P on the half-life of the RNase Y substrate *rpsO* mRNA. More important, SR7P had a repercussion on survival of *B. subtilis* cells after stress: after ethanol stress, SR7P increased survival about fourfold, after heat and acid stress, about 1.5- to twofold (Fig. 10). These biological effects might be explained by an influence of SR7P on the degradation of a number of RNase Y substrates, which in turn affects targets in different pathways.

The discovery of an impact of two 39 aa proteins, SR1P and SR7P, on components of the putative *B. subtilis* degradosome suggests that sproteins might also play a broader role in fine-tuning RNA degradation in other bacteria. Like SR1P [32], SR7P is restricted to the Bacillales (Fig. 11, Fig. S5). Sequence homology could not be found to any other sprotein in other Gram-positive or in Gram-negative bacteria. However, it is not excluded that in degradosomes of other bacteria that contain enolase as scaffolding component [51], small proteins with a different primary sequence play a comparable role under specific stress conditions.

The SigB-dependent RNA encoding SR7P was formerly reported as S1136, an antisense RNA whose convergent transcription to *rpsD* mRNA results in reduced amounts of the small ribosomal subunit under ethanol stress [37]. We found that this RNA, which we renamed as SR7, has in addition to its antisense RNA function an mRNA function: It encodes SR7P which – via enolase binding – affects RNA degradation and cell survival. Therefore, SR7 is after SR1 [29,30,33,38] the second dual-function regulatory RNA identified in *Bacillus subtilis*. All dual-function regulatory RNAs reported so far act only *in trans* on their target mRNAs [24]. By contrast, Mars *et al*. found that SR7 acts only *in cis* – most probably due transcriptional interference [37]. Consequently, the class of dual-function regulatory RNAs can be extended by dual-function antisense RNAs.

Future investigations will focus on the structure of SR7P, structural alterations of both interacting partners, SR7P and enolase, upon binding, and on mapping of the enolase/SR7P interaction surface, as initiated already [47]. Moreover, transcriptomics will show whether the abundance of other RNase Y substrates is also affected by SR7P. Eventually, we aim to elucidate how SR7P affects cell survival under different stress conditions.

## Materials and Methods

### Strains, media and growth conditions

*E. coli* strains DH5α and BL21DE3 and *B. subtilis* strains DB104 [52] were used. TY medium served as complex medium for *E. coli* [30]. All *B. subtilis* strains were grown in TY medium until OD_560_ = 4.

### Enzymes and chemicals

Chemicals used were of the highest purity available. Q5 DNA polymerase, T7 RNA polymerase, CIP and polynucleotide kinase were purchased from NEB, Firepol Taq polymerase from Solis Biodyne, and sequenase from Affymetrix.

### Isolation of chromosomal DNA from Bacillus subtilis

0.5 ml of *B. subtilis* DB104 stationary phase culture was centrifuged, the pellet washed with 1 ml TES (10 mM Tris-HCl pH 8.0, 1 mM EDTA, 100 mM NaCl), re-suspended in 750 µl TES containing 25 µl lysozyme (10 mg/ml) and incubated at 37 °C for 5 min. 50 µl pronase E (20 g/l) were added and mixed gently. Subsequently, 50 µl 10% SDS were added, mixed gently, followed by a 30 min incubation at 37 °C. Afterwards, one phenol/chloroform extraction and one chloroform extraction were performed using decapitated 1 ml tips, and the final supernatant was added to 2 ml 96 % ethanol. After one min of gentle shaking the precipitated DNA was carefully taken out with a yellow tip, dissolved in 100 µl bidest, incubated at 37 °C for 30 min followed by one hour on ice.

### Primer extension

Primer extension experiments were carried out as described [29] using total RNA from *B. subtilis* strain DB104, DB104 (*Δrny::spec*^*R*^) and DB104 (*Δrnc::cm*^*R*^) and 5’-labelled primer SB3182 (all primers are listed in Table S1). Primer extension experiments were carried out as described [29]. As control served a sequencing reaction with primer SB1171 using pUC19-*sr7*_Pro_ as template.

### *In vitro* transcription, preparation of total RNA and Northern blotting

*In vitro* transcription was performed as described [38]. Preparation of total RNA and Northern blotting including the determination of RNA half-lives were carried out as described previously [29] except that 1 ml time samples were taken and directly added to 250 μl RNAprotect Bacteria Reagent (Qiagen) or to 250 μl containing 5 % phenol and 95 % ethanol, vortexed and incubated for 5 min at room temperature. After centrifugation, the pellets were flash-frozen in liquid nitrogen and stored until use at −20 °C.

### Strain constructions

All PCRs were performed with Q5 polymerase using 25 cycles with an annealing temperature of 48 °C. If not stated otherwise, primers (listed in Table S1) and as template, chromosomal DNA of *B. subtilis* DB104 were used. All constructed recombinant strains were confirmed by sequencing using the primers listed in Table S1.

### sr7p_C-FLAG_ and sr7p_C-His10_ knock-in strains

A 340 bp fragment encoding SR7P_C-FLAG_ was obtained in a two-step PCR. In PCR 1 with primer pair SB2963/SB2978, a C-terminal FLAG-tag was added at the end of the *sr7* ORF. In the subsequent PCR with primer SB2843, a stop codon was introduced after the FLAG-tag sequence and the native terminator replaced by the heterologous BsrF terminator [39] yielding the *sr7p*_*C-FLAG*_ fragment. The chloramphenicol resistance (cm^R^) cassette was amplified from plasmid pINT12C [46] with primer pair SB2938/SB2939. This cassette has 20 bp complementarity to the *sr7p*_*C-FLAG*_ fragment on its 5’ end and 20 bp complementarity to the back cassette on its 3’ end. A 1 kb front cassette that has 20 bp complementarity on its 3’ end to the *sr7p*_*C-FLAG*_ fragment was obtained using primer pair SB2964/SB2966. Similarly, a 1 kb back cassette was generated with primer pair SB2967/SB2968 which has a 5’ 20 bp complementarity to the cm^R^ gene. The four PCR fragments were purified and joined by a 10-cycles’ primer-less PCR. Afterwards, the final 3 kb fragment was produced using a 25 cycles’ PCR with primer pair SB2964/SB2968. This fragment was used to transform *B. subtilis* DB104. The resulting strain was designated DBSR7PF (all strains are listed in Table 1).

**Table 1:** Bacterial strains used in this study.

| Strain | Genotype | Reference |
| --- | --- | --- |
| *E. coli* DH5α | *fhu2*, Δ(*argF-lacZ*), *U169*, *phoA*, *glnV44*, Φ80, Δ(*lacZ*)*M15*, *gyrA96*, *recA1*, *relA1*, *endA1*, *thi-1*, *hsdR17* | [54] |
| *E. coli* SSC420 | overexpression strain for *B. subtilis* *rny<sub>C-His6</sub>* without N-terminal transmembrane domain | Harald Putzer, Paris |
| *E. coli* BL21DE3 | *fhuA2* [*lon*] *ompT* *gal* (λ *DE3*) [*dcm*] Δ*hsdS*<br>λ *DE3* = λ *sBamHlo* Δ*EcoRI-B*<br>*int::(lacI::PlacUV5::T7 gene1)* *i21* Δ*nin5* | NEB |
| *B. subtilis* GP1215 | *B. subtilis* 168 with *eno<sub>N-Strep</sub>* at native locus | Jörg Stülke, Göttingen |
| *B. subtilis* BKK33900 | *trpC2*, *eno::kan<sup>R</sup>* | Byoung-Mo Koo, San Francisco |
| *B. subtilis* DB104 | *His*, *nprR*, 2 *nprE18*, Δ*aprA3* | [51] |
| DB104(*amyE::gapA<sub>Strep</sub>*) | *his nprR2 nprE18 ΔaprA3*, Δ*amyE::pMGG19* | [33] |
| *B. subtilis* BKK14990 | *B. subtilis* 168 lacking the *ylbF* gene, Km<sup>R</sup> | Ohio strain collection |
| *B. subtilis* DB(Δ*ylbF*) | *B. subtilis* DB104 lacking the *ylbF* gene, Km<sup>R</sup> | This study |
| *B. subtilis* DBE(Δ*ylbF*) | *B. subtilis* DB104 with *eno-N-Strep*, Δ*ylbF*, Spec<sup>R</sup>, Km<sup>R</sup> | This study |
| *B. subtilis* DBSR7PF | *B. subtilis* DB104 with *sr7p<sub>C-FLAG</sub>*, Cm<sup>R</sup> | This study |
| *B. subtilis* DBSR7PFE | *B. subtilis* DB104 with *sr7p<sub>C-FLAG</sub>* and *eno<sub>N-Strep</sub>*, Spec<sup>R</sup>, Cm<sup>R</sup> | This study |
| *B. subtilis* DBSR7PFEΔ*ylbF* | *B. subtilis* DBSR7PFE with Δ*ylbF*, Spec<sup>R</sup>, Cm<sup>R</sup>, Km<sup>R</sup> | This study |
| *B. subtilis* DBSR7PH | *B. subtilis* DB104 with *sr7p<sub>C-His</sub>* Cm<sup>R</sup> | This study |
| *B. subtilis* DBSR7PHE | *B. subtilis* DB104 with *sr7p<sub>C-His10</sub>* and *eno<sub>N-Strep</sub>*, Spec<sup>R</sup>, Cm<sup>R</sup> | This study |
| *B. subtilis* DB104(Δ*p<sub>sr7</sub>*) | *B. subtilis* DB104 lacking only *p<sub>sr7</sub>*, Spec<sup>R</sup> | This study |
| *B. subtilis* DB104(Δ*sr7*) | DB104 with a deletion of the entire *sr7* gene, Cm<sup>R</sup> | This study |
| *B. subtilis* DBE(Δ*sr7*) | DB104 with a deletion of the entire *sr7* gene with *eno<sub>N-Strep</sub>*, Spec<sup>R</sup>, Cm<sup>R</sup> | This study |
| *B. subtilis* DBSR7P-1 | DB104 with *sr7* carrying a start- to stop codon mutation | This study |
| *B. subtilis* DBPF<sub>C-His</sub> | DBSR7PF with *pfkA<sub>C-His6</sub>* at its native locus, Cm<sup>R</sup>, Km<sup>R</sup> | This study |
| *B. subtilis* DBPF<sub>C-FLAG</sub> | DBSR7PF with *pfkA<sub>C-FLAG</sub>* at its native locus, Cm<sup>R</sup>, Km<sup>R</sup> | This study |
| *B. subtilis* DB104 (Δ*rny*) | DB104 with a replacement of the *rny* by the Spec<sup>R</sup> gene | [46] |
| *B. subtilis* DB104 (Δ*rnc*) | DB104 with replacement of the *rnc* by the Cm<sup>R</sup> gene | [31] |
| *B. subtilis* DB2184F | *B. subtilis* DB104 with *sp2184<sub>C-FLAG</sub>*, Cm<sup>R</sup> | Müller, unpublished |
| DB104 (*amyE::sr7p<sub>C-FLAG</sub>*) | DB104 with IPTG inducible *sr7p<sub>C-FLAG</sub>* in *amyE* locus, Spec<sup>R</sup> | This study |

A 1 kb fragment with 10 histidine residues at the 3’ end of the *sr7* ORF was obtained using primer pair SB3013/SB3575. This fragment has 20 bp complementarity to the 5’ primer region of the cm^R^ (see above) gene. A 1 kb back cassette with 20 bp complementarity to the 3’ region of the cm^R^ gene was generated using primer pair SB2967/SB3156. All three fragments were joined, primer pair SB3013/SB3156 added, a final 3 kb fragment was amplified and used to transform *B. subtilis* DB104 yielding strain DBSR7PH.

### Construction of the *sr7* promoter knock-out strain, the *Δsr7* strain and the start- to stop-codon mutant strain

To delete the *sr7* promoter, a 1 kb front cassette upstream of the *sr7* gene was generated by PCR 1 with primer pair SB3013/ SB3014. The 0.8 kb spectinomycin resistance (spec^R^) gene was obtained by PCR 2 on plasmid pMG16 [46] using primer pair SB3039/SB3040. The 3’ end of the front cassette has a 20 bp complementarity to the 5’ region of the spec^R^ gene. The 1.14 kb back cassette comprising the promoterless *sr7* gene together with the *rpsD* gene located downstream on the complementary strand was generated by PCR 3 with primer pair SB3042/SB3020. It displays a 20 bp complementarity to the 3’ end of the spec^R^ gene. As above, the purified fragments were joined, a 3 kb fragment obtained with primer pair SB3013/ SB3020 and used to transform *B. subtilis* DB104 resulting in promoter knockout strain DBSR7PΔP.

To delete the *sr7* ORF, a 1 kb front cassette was generated using primer pair SB3013/SB3014 with 20 bp complementarity to the 5’ of the *cm*^*R*^ cassette. The 1 kb back cassette was amplified using primer pair SB3243/SB3156. The BsrF terminator [39] was added to the *rpsD* ORF present on the opposite strand with 20 bp complementarity to the 3’ end of the *cm*^*R*^ gene. The fragments were joined, a final 3 kb fragment obtained as above using primer pair SB3013/SB3156 and used to transform *B. subtilis* DB104 resulting in DB104 *(Δsr7*).

To construct an *sr7* start- to stop-codon mutant strain, a 1 kb fragment upstream of the *sr7* locus was amplified using primer pair SB3013/SB3014. This fragment has 20 bp complementarity to the 5’ region of the *cm*^*R*^ gene. The *sr7* start codon was replaced by a stop codon through a PCR with primer pair SB3017/SB3227 introducing simultaneously a 20 bp complementarity to the 3’ region of the *cm*^*R*^ gene. A 1 kb back fragment was generated with primer pair SB3226/SB3156. All four fragments were joined, a final 3.2 kb fragment was obtained SB3013/SB3156 and used to transform *B. subtilis* DB104 yielding strain DBSR7P-1.

### pfkA_C-His6_ *and* pfkA_C-FLAG_ *knock-in strains*

To add a His or a FLAG tag to the C-terminus of the *pfkA* ORF, a 1 kb front cassette encoding 6 His residues or 3x FLAG was generated using primer pair SB3269/SB3347 or SB3269/SB3270, respectively. The neomycin resistance (km^R^) resistance gene was amplified from plasmid pMG9 using primer pair SB3039/SB3180. In the subsequent PCR with primer SB3267 or SB3167 a stop codon and a 20 bp complementarity to the 5’ region of the *km*^*R*^ cassette were introduced. A 1 kb back cassette with 20 bp complementarity to the 3’ primer region of the *km*^*R*^ gene was obtained using primer pair SB3271/SB3272. All fragments were purified, joined in a final PCR with SB3269/SB3272 as above to generate a 3 kb fragment which was used to transform *B. subtilis* DBSR7PF yielding *B. subtilis* DBPF_C-His_ or DBPF_C-FLAG_.

### Plasmid constructions

For the construction of a translational *ncr2360*-*lacZ* fusion (later renamed as *sr7p-lacZ* fusion) under control of the constitutive heterologous promoter pIII [35] a PCR fragment was generated with primer pair SB2816/SB2817, cleaved with BamHI and EcoRI and inserted into the BamHI/EcoRI pGAB1 vector resulting in pGABP2360. For the construction of a plasmid for IPTG-inducible overexpression SR7P_C-FLAG_ a PCR fragment lacking the *sr7* promoter was obtained on chromosomal DNA using primer pair SB2872/SB2878. In the subsequent PCR, the native stop codon was replaced by a 3 x FLAG tag with primer pair SB2872/SB3439. The resulting fragment was cleaved with Hind III and SphI and inserted into the pDR111 [53] Hind III/SphI vector. Both recombinant vectors pGAB2360 and pDRSR7P were amplified in *E. coli* DH5α, linearized with ScaI and integrated into the *amyE* locus of *B. subtilis* strains DB104 and DB104 *(Δsr7*), respectively.

### Preparation of protein crude extracts for the detection of SR7P_C-FLAG_, Western Blotting and Far-Western Blotting

Protein crude extracts were prepared by either sonication (see below) or DNase I/lysozyme treatment. In the latter case, pellets were treated with 10 mg/ml lysozyme/1 mg/ml DNase I in PBS for 30 min at 37 °C followed by two 5 min centrifugation steps (10 min at RT) to obtain the supernatant.

Western blotting for the detection of SR7P_C-FLAG_ was performed as follows: The pellet from a 100 ml culture grown in TY was resolved in 4 ml TBS and sonicated three times for 5 min. Supernatants obtained by two 10 min centrifugation steps at 4 °C were run through an anti-FLAG M2 column (bed volume 200 µl), washed 6 times in 200 µl TBS and followed by 5 elution steps with 200 μl of 100 mM glycine HCl pH 3.5. 1 ml of elution fractions was mixed with 1 ml 50 % ice-cold TCA, placed on ice for 30 min and centrifuged for 30 min at 13000 rpm. The pellet was washed three times with ice-cold acetone (20 min centrifugation), air-dried for 5 min and dissolved in 50 μl 2x Laemmli buffer. 10 μl were loaded onto a 15 % SDS PAA gel run at 35 mA for 75 min and transferred onto PVDF membranes at 12 V for 40 min by semidry blotting in transfer buffer (5.8 g Tris-HCl, 2.9 g glycine, 0.37 g SDS and 200 ml methanol per liter). Membranes were blocked for 1 h in PBST [31] with 0.5% gelatine and incubated for 1 h with M2 anti-FLAG antibodies (1:2500) followed by 5 times 8 min washing in PBST. Subsequently, membranes were incubated with secondary antibodies (horseradish peroxidase conjugated anti-mouse) for 1 h followed by 5 washing steps. For development, membranes were incubated in 20 ml substrate solution (50 mM Tris pH 7.5, 0.9 g/ml diaminobenzidine and 20 µl H_2_O_2_) until bands were visible. The reaction was stopped by washing in distilled water. For enolase detection, rabbit against enolase antibodies (1:5000) were used as primary and horseradish peroxidase conjugated anti-rabbit antibodies (1:2500) as secondary antibodies.

Detection by chemiluminescence was used for RNase Y, PfkA_C-His6_, quantification of SR7P amounts and RNase Y after crude extracts were treated with RNase A: The membrane was washed briefly with deionized water and the pH adjusted with a solution containing 100 mM NaCl and 100 mM Tris-HCl pH 9.5. Afterwards, the membrane was incubated with developing solution (Invitrogen Novex AP Chemiluminescent substrate) for 5 min in the dark and signals were detected with a CCD camera (ImageQuant LAS 4000).

For Far-Western blotting, proteins were separated on 10 % or 15 % SDS/PAA gels and transferred onto PVDF membranes and afterwards blocked for 2 h. Blots were incubated overnight with 10 ml PBST-gelatine or PBST-gelatine supplemented with either 100 μg Eno_N-Strep_ purified from wild-type or *Δsr7 B. subtilis* strains or 170 μg RNase Y_C_-_His6_ purified from *E. coli*. Binding of Eno_N-Strep_ and RNase Y_C_-_His6_ was detected by incubation with mouse-anti-Strep-tag antibody (1:1000; IBA Göttingen) or mouse-anti-His-tag antibody (1:2000; IBA Göttingen), respectively, and subsequently alkaline phosphatase coupled anti-mouse antibody (1:7500; Santa Cruz Biotechnology).

### Purification of FLAG-tagged, Strep-tagged and His-tagged proteins from *B. subtilis*

*B. subtilis* strain DBSR7PF was grown in 0.8 l TY medium till onset of stationary phase and harvested by centrifugation at 4 °C and 8000 rpm. Pellets were frozen overnight. Subsequently, they were disrupted in 2 ml total volume of equilibration buffer (150 mM NaCl, 100 mM Tris-HCl pH 8.0, 1 mM EDTA) for 1.5 min at 2000 rpm in the Mikro-Dismembrator (Sartorius). Resulting crude extracts were subjected to sonication in 30 ml buffer and afterwards centrifuged twice at 13.000 rpm and 4 °C. Supernatants were applied to M2 anti-FLAG column (strain DBSR7PF) or to Strep-Tactin columns (strain GP1215). Washing, elution and regeneration of the 1 ml columns were performed according to the manufacturers’ instructions. Six 500 μl elution fractions were collected for each column, and 25 μl of each fraction tested on SDS-PAA gels. The purification of SR7P_C-His10_ was performed as described before [31].

### RNA degradation assay

Two μl *in vitro* transcribed, [^32^P]-γ ATP labelled RNA (10.000 cpm) were incubated with 1 μl 10x reaction buffer (200 mM Tris-HCl pH 8.0; 80 mM MgCl_2_, 1 M NH_4_Cl; 0.5 mM DTT including 1 U RNasin) and 7 μl diluted protein for 30 min at 37 °C. The reaction was stopped by addition of 10 μl formamide loading dye (Licht et al., 2005) and 5 min at 95 °C. Samples were separated on 6 % denaturing PAA gels. Dried gels were analysed by phosphorImaging using Aida Image Analyzer v.4.5.

### Survival assay

Cultures were grown in TY medium at 37 °C to OD_560_=4, fivefold diluted in either fresh TY medium or fresh TY with final concentrations of 12 % ethanol, 4 M NaCl, 15 mM MnCl_2_ or HCl (final pH: 4.0). Growth was continued for 30 min under stress conditions. For heat stress, cultures were shifted to 57 °C for 10 min. Subsequently, serial dilutions were plated on TY agar plates, incubated overnight at 37 °C and afterwards, colonies counted.

The control culture was diluted and plated immediately without 30 min additional incubation to avoid further growth. All stress conditions were tested preliminarily and optimized for a significant reduction of survival after 30 min or (heat stress) 10 min. Therefore, the stressor concentrations used were much higher than those required for induction of the SigB-dependent promoter p_*sr7*_.

## Abbreviations

aa: amino acid
nt: nucleotide
bp: base pair
ORF: open reading frame
PAA: polyacrylamide
sRNA: small RNA
RBS: ribosome binding site
sprotein: small protein.

## Acknowledgements

The authors thank Bernhard Schlott (FLI Jena) for performing the mass spectrometry analysis to identify enolase. We thank Jörg Stülke (Göttingen) for strain GP1215, antibodies against native RNase Y and comments on the manuscript. We are grateful to Harald Putzer (Paris) for *E. coli* strain SSC420 for overexpression of RNase Y_C-His6_ and to Byoung-Mo Koo (San Francisco) for the *Δeno* strain. Furthermore, we thank Laura Remmel and Nina Kubatova (AG Schwalbe, Frankfurt/Main) for SR7P purified from *E. coli*.

## References

1. Hobbs EC, Fontaine F, Yin X, Storz G. An expanding universe of small proteins. Curr Opin Microbiol. 2011;14:167–173.

2. Storz G, Wolf YI, Ramamurthi KS. Small proteins can no longer be ignored. Annu Rev Biochem. 2014; 83:753–777.

3. Orr MW, Mao Y, Storz G, Quian SB. Alternative ORFS and small ORFs: shedding light on the dark proteome. Nucleic Acids Res. 2020;48:1029–1042.

4. Andrews SJ, Rothnagel JA. Emerging evidence for functional peptides encoded by short open reading frames. Nat Rev Genet. 2014;15:193–204.

5. Brantl S, Jahn N. sRNAs in bacterial type I and type III toxin-antitoxin systems. FEMS Microbiol Rev. 2015;39:413–427.

6. Jahn N, Brantl S, Strahl H. Against the mainstream: the membrane-associated type I toxin BsrG from *Bacillus subtilis* interferes with cell envelope biosynthesis without increasing membrane permeability. Mol Microbiol. 2015;98:651–666.

7. Vogel J, Luisi BF. Hfq and its constellation of RNA. Nat Rev Microbiol. 2011;9:578–589.

8. Kavita K, de Mets F, Gottesman S. New aspects of RNA-based regulation by Hfq and its partner sRNAs. Curr Opin Microbiol. 2018;42:53–61.

9. Vakulskas CA, Potts AH, Babitzke P, Ahmer BMM, Romeo T. Regulation of bacterial virulence by Csr (Rsm) systems. Microbiol. and Molecular Biol Rev. 2015;79:193–224.

10. Müller P, Gimpel M, Wildenhain T, Brantl S. A new role for CsrA: promotion of complex formation between an sRNA and its mRNA target in *Bacillus subtilis*. RNA Biol. 2019;16:972–987.

11. Gaballa A, Antelmann H, Aguilar C, Khakh SK, Song KB, Smaldone T, Helmann JD. The *Bacillus subtilis* iron-sparing response is mediated by a Fur-regulated small RNA and three small, basic proteins. Proc Natl Acad Sci USA. 2008;105:11927–11932.

12. Smaldone GT, Revelles O, Gaballa A, Sauer U, Antelmann H, Helmann JD. A global investigation of the *Bacillus subtilis* iron-sparing response identifies major changes in metabolism. J Bacteriol. 2012; 194:2594–2605.

13. Martin JE, Waters LS, Storz G. The *E. coli* small protein MntS and exporter MntP optimize the intracellular concentration of manganese. PLoS Genet. 2015;11:e1004977.

14. Cutting S, Anderson M, Lysenko E, Page A, Tomoyasu T, Tatematsu K, Ogura T. SpoVM, a small protein essential to development in *Bacillus subtilis,* interacts with the ATP-dependent protease FtsH. J Bacteriol.1997;17:5534–5542.

15. Gill RL, Castaing JP, Hsin J, Tan IS, Wang X, Huang KC et al.Structural basis for the geometry-driven localization of a small protein. Proc Natl Acad Sci USA. 2015;112:E1908–15.

16. Ramamurthi KS, Lecuyer S, Stone HA, Losick R. Geometric cue for protein localization in a bacterium. Science. 2009;323:1354–1357.

17. Cunningham KA, Burkholder WF. The histidine kinase inhibitor Sda binds near the site of autophosphorylation and may sterically hinder autophosphorylation and phosphotransfer to Spo0F. Mol Microbiol. 2009;71:659–677.

18. Ray S, Kumar A, Panda D. GTP regulates the interaction between MciZ and FtsZ: A possible role of MciZ in bacterial cell division. Biochemistry 2013;52:392–401.

19. Bisson-Filho AW, Discola KF, Castellen P, Blasios V, Martins A, Sforça ML, Gueiros-Filho FJ. FtsZ filament capping by MciZ, a developmental regulator of bacterial division. Proc Natl Acad Sci USA. 2015;12:2130–2138.

20. Brantl S. Bacterial chromosome-encoded small regulatory RNAs. Future Microbiol. 2009;4: 85–103.

21. Brantl S. Acting antisense: plasmid- and chromosome-encoded sRNAs from Gram-positive bacteria. Future Microbiol. 2012;7:853–871.

22. Brantl S, Brückner R. Small regulatory RNAs from low-GC Gram-positive bacteria. RNA Biol. 2014;11:443–456

23. Wagner EGH, Romby P. Small RNAs in bacteria and archaea: who they are, what they do, and how they do it. Adv Genet. 2015;90:133–208.

24. Gimpel M, Brantl S. Dual function small regulatory RNAs in bacteria. Mol Microbiol. 2017;103: 387–397.

25. Wadler CS, Vanderpool CK. A dual function for a bacterial small RNA: SgrS performs base pairing dependent regulation and encodes a functional polypeptide. Proc Natl Acad Sci USA. 2007;104:20454–20459.

26. Bronesky D, Wu Z, Marzi S, Walter P, Geissmann T, Moreau K, Vandenesch F, Caldelari I, Romby P. *Staphylococcus aureus* RNAIII and its regulon link quorum sensing, stress responses, metabolic adaptation, and regulation of virulence gene expression. Annu Rev Microbiol. 2016;8:299–316.

27. Kaito C, Saito Y, Ikuo M, Omae Y, Mao H, Nagano G. et al. Mobile genetic element SCCmec-encoded psm-mec RNA suppresses translation of *agrA* and attenuates MRSA virulence. PLoS Pathog. 2013; 9:e1003269.

28. Mangold M, Siller M, Roppenser B, Vlaminckx BJ, Penfound TA, Klein R, Novak R, Novick RP, Charpentier E. Synthesis of group A streptococcal virulence factors is controlled by a regulatory RNA molecule. Mol Microbiol. 2004;53:1515–1527.

29. Licht A, Preis S, Brantl S. Implication of CcpN in the regulation of a novel untranslated RNA (SR1) in *B. subtilis*. Mol Microbiol. 2005;58:189–206.

30. Heidrich N, Chinali A, Gerth U, Brantl S. The small untranslated RNA SR1 from the *B. subtilis* genome is involved in the regulation of arginine catabolism. Mol Microbiol. 2006;62:520–536.

31. Gimpel M, Heidrich N, Mäder U, Krügel H, Brantl S. A dual function sRNA from *B. subtilis*: SR1 acts as a peptide encoding mRNA on the *gapA* operon. Mol Microbiol. 2010;76: 990–1009.

32. Gimpel M, Preis H, Barth E, Gramzow L, Brantl S. SR1 – a small RNA with two remarkably conserved functions. Nucleic Acids Res. 2012;40:11659–11672.

33. Gimpel M, Brantl S. Dual-function sRNA-encoded peptide SR1P modulates moonlighting activity of *B. subtilis* GapA. RNA Biol. 2016;13:916–926.

34. Irnov I, Sharma CM, Vogel J, Winkler WC. Identification of regulatory RNAs in *Bacillus subtilis*. Nucleic Acids Res. 2010;38:6637–6651.

35. Brantl S, Nuez B, Behnke D. *In vitro* and *in vivo* analysis of transcription within the replication region of plasmid pIP501. Mol Gen Genet. 1992;234:105–112.

36. Commichau FM, Rothe FM, Herzberg C, Wagner E, Hellwig D, Lehnik-Habrink M, et al.Novel activities of glycolytic enzymes in *Bacillus subtilis*: interactions with essential proteins involved in mRNA processing. Mol Cell Proteomics. 2009;8:1350–1360.

37. Mars, RA, Mendonça K, Denham EL, van Dijl JM. The reduction in small ribosomal subunit abundance in ethanol-stressed cells of *Bacillus subtilis* is mediated by a SigB-dependent antisense RNA. Biochim Biophys Acta. 2015;1853:2553–2559.

38. Heidrich N, Moll I, Brantl S. *In vitro* analysis of the interaction between the small RNA SR1 and its primary target *ahrC* mRNA. Nucleic Acids Res. 2007;35:331–346.

39. Preis H, Eckart RA, Gudipati RK, Heidrich N, Brantl S. CodY activates transcription of a small RNA in *Bacillus subtilis*. J Bacteriol. 2009;191:5446–5457.

40. Newman JA, Hewitt L, Rodrigues C, Solovyova A, Harwood CR, Lewis RJ. Dissection of the network of interactions that links RNA processing with glycolysis in the *Bacillus subtilis* degradosome. J Mol Biol. 2012;416:121–136.

41. Lehnik-Habrink M, Schaffer M, Mäder U, Diethmaier C, Herzberg C, Stülke J. RNA processing in *Bacillus subtilis*: identification of targets of the essential RNase Y. Mol Microbiol. 2011;81:1459–1473.

42. Shahbabian K, Jamalli A, Zig L, Putzer H. RNase Y, a novel endoribonuclease, initiates riboswitch turnover in *Bacillus subtilis*. EMBO J. 2009;28:3523–2533.

43. Yao S, Bechhofer D. Initiation of decay of *Bacillus subtilis rpsO* mRNA by endoribonuclease RNase Y. J Bacteriol. 2010;192:3279–3286.

44. DeLoughery A, Lalanne JB, Losick R, Li GW. Maturation of polycistronic mRNAs by the endoribonuclease RNase Y and its associated Y-complex in *Bacillus subtilis.* Proc Natl Acad Sci USA. 2018;115:E5585–E5594.

45. Jahn N, Preis H, Wiedemann C, Brantl S. BsrG/SR4 from *Bacillus subtilis* – the first temperature-dependent type I toxin-antitoxin system. Mol Microbiol. 2012;83:579–598.

46. Müller P, Jahn N, Ring C, Maiwald C, Neubert R, Meißner C, Brantl S. A multistress responsive type I toxin-antitoxin system: *bsrE/SR5* from *Bacillus subtilis*. RNA Biol. 2016;13:511–523.

47. Kubatova N, Pyper DJ, Jonker HRA, Saxena K, Remmel L, Richter C, Brantl S, Evguenieva-Hackenberg E, Hess WR, Klug G, Marchfelder A, Soppa J, Streit W, Mayzel M, Orekhov VY, Fuxreiter M, Schmitz RA, Schwalbe H. Rapid biophysical characterization and NMR spectroscopy structural analysis of small proteins from Bacteria and Archaea. Chembiochem. 2019;doi 10.1002/cbic.201900677.

48. Lehnik-Habrink M, Lewis RJ, Mäder U, Stülke J. RNA degradation in *Bacillus subtilis*: an interplay of essential endo- and exoribonucleases. Mol Microbiol. 2012;84:1005–1017.

49. Yang J, Gao M, Xiong J, Su Z, Huang Y. Features of molecular recognition of intrinsically disordered proteins via coupled folding and binding. Protein Science. 2019;28:1952–1965.

50. Nurmohamed S, Vincent HA, Titman CM, Chandran V, Pears MR, D. D, Griffin JHL, Callaghan AJ, Luisi BF. Polynucleotide phosphorylase activity may be modulated by metabolites in *Escherichia coli.* J Biol Chem. 2011;286:14315–14323.

51. Carpousis AJ. The RNA degradosome of *Escherichia coli*: an mRNA-degrading machine assembled on RNase E. Annu Rev Microbiol. 2007;61:71–87.

52. Kawamura F, Doi RH. Construction of a *Bacillus subtilis* double mutant deficient in extracellular alkaline and neutral proteases. J Bacteriol. 1984;160:442–444.

53. van Ooij C, Losick R. Subcellular localization of a small sporulation protein in *Bacillus subtilis*. J Bacteriol. 2003;185:1391–1398.

54. Hanahan D. Studies on transformation of *Escherichia coli* with plasmids. J Mol Biol. 1983;166:557–580.

