## Supplementary Tables and Figures for "SR7 - a dual function antisense RNA from *Bacillus subtilis*"

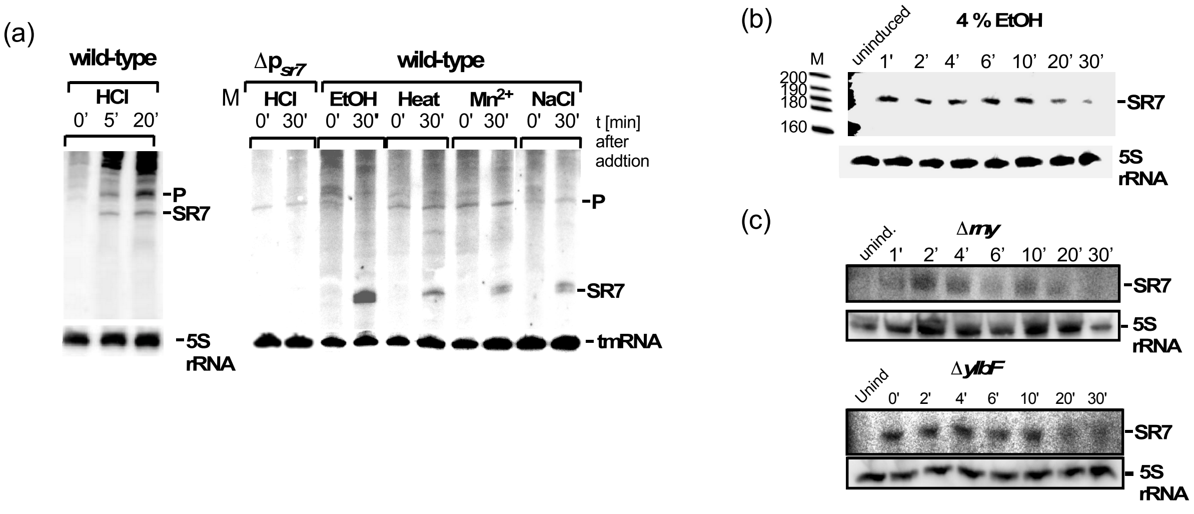


**Fig. S1. Transcription of the SigB-dependent SR7 is induced under five different stress conditions**

*B. subtilis* DB104 was grown in TY medium until stationary phase at 37 °C and subjected to stress (0.5 M NaCl, 4 % ethanol, 1 mM $Mn^{2+}$, pH 5.0 or 48 °C) for 15 min. Total RNA was prepared from flash-frozen samples, separated on 6 % denaturing PAA gels and blotted onto nylon membranes.

(a) For comparison, an isogenic strain carrying a deletion of the SigB promoter upstream of the *sr7* gene was used. Samples on the left side were separated in a 6 % denaturing PAA gel, on the right side on a 4 % denaturing PAA gel. SR7 was detected by hybridization with a [$^{32}$P]-αUTP-labelled riboprobe. Autoradiograms of the Northern blots are shown. Loading errors were corrected by reprobing with a [$^{32}$P] γATP-labelled oligonucleotide specific for 5S rRNA or by reprobing with a [$^{32}$P]-αUTP-labelled riboprobe against tmRNA. P, processing products from a longer RNA originating at the constitutive *tyrS* promoter.

(b) Northern blots of DB104 RNA isolated after induction with 4 % ethanol for 15 min, followed by addition of rifampicin to prevent further transcription initiation. It can be seen that SR7 is not less stable than in after induction with 0.5 M NaCl (Fig. 2)

(c) Northern blot RNA isolated from DB104(Δ*rny*) and DB104(Δ*ylbF*) after salt stress (0.5 M NaCl) for 30 min.

Figure S1 Ul Haq *et al.*

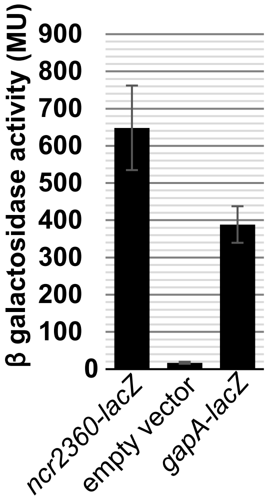


**Fig. S2. The putative SD sequence of *ncr2360* (later renamed as *sr7p*) is recognized in *B. subtilis.*** A translational *ncr2360-lacZ* fusion comprising the putative *ncr2360* SD and the first three codons was constructed under control of constitutive heterologous promoter pIII using vector pGAB1 and integrated into the *amyE* locus of *B. subtilis* strain DB104. Cultures were grown in complex TY medium until $OD5_{60}$ = 4 and used for the measurement of β- galactosidase activities as described (Gimpel *et al*., 2010). The empty vector pGAB1 served as negative control and pGAB1 carrying a translational *gapA-lacZ* fusion served as positive control.

Fig. S2 Ul Haq *et al.*

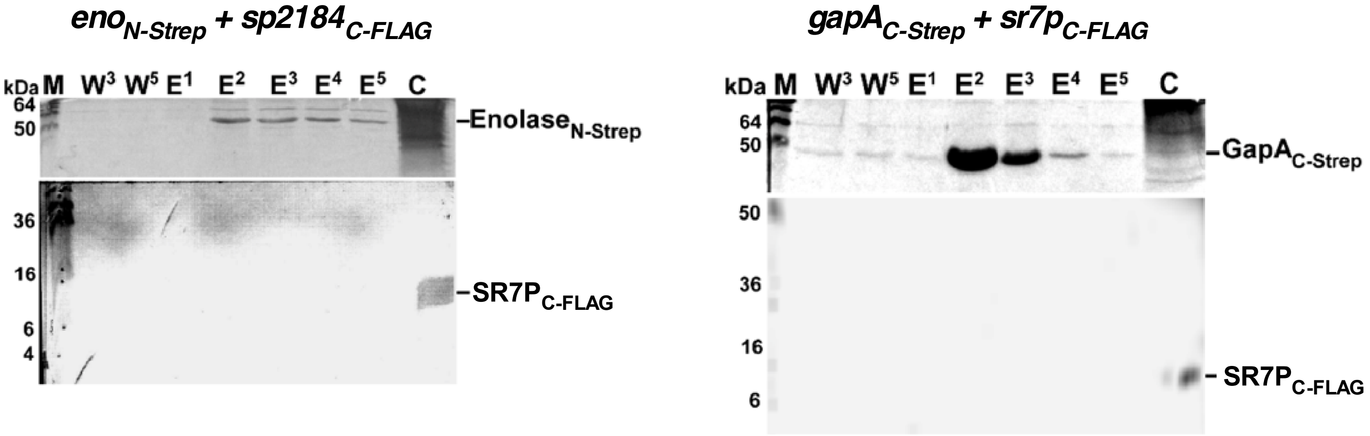


**Fig. S3. Exclusion of an interaction of Eno$_{N\text{-}Strep}$ with any FLAG-tag and of SR7P$_{C\text{-}FLAG}$ with any Strep-tag.**
*B. subtilis* strain DB104 with chromosome-encoded Eno$_{N\text{-}Strep}$ and sprotein SP2184$_{C\text{-}FLAG}$ (left) and *B. subtilis* strain DB104 with chromosome-encoded GapA$_{C\text{-}Strep}$ and SR7P$_{C\text{-}FLAG}$ (right) were grown in complex TY medium until $OD_{560}$ = 4.5., crude protein extracts prepared as described in *Materials and Methods* and passed through Streptactin columns. Washing fractions 3 and 5 as well as 5 elution fractions were separated on 15 % SDS PAA gels, which were stained with Coomassie blue (above). Afterwards, gels were subjected to Western blotting with anti-FLAG antibodies (see below). As positive controls (C) SRP7$_{C\text{-}FLAG}$ was loaded on each gel. No SP2184 was detectable on the left gel and no SR7P$_{C\text{-}FLAG}$ on the right gel in the elution fractions.

Figure S3 Ul Haq *et al.*

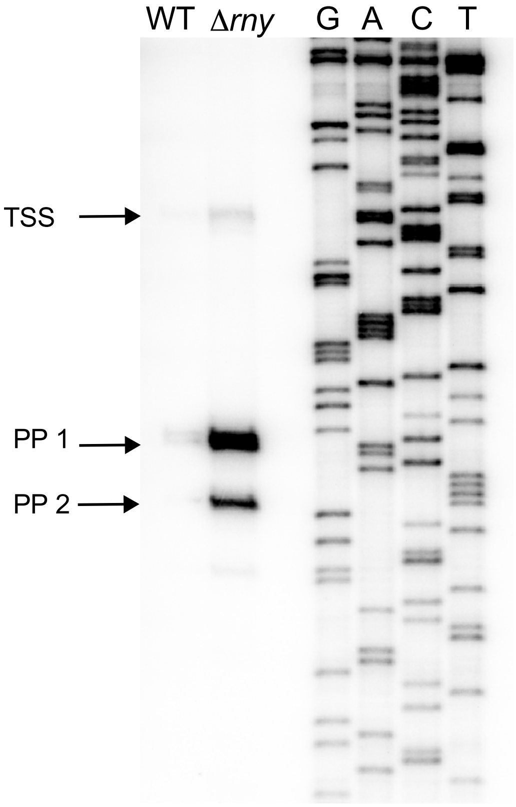


-35 box -10 box

5' **TTGCAA**CGCACGCAAATTTTATTC**TAAAAT**ATTTGCATATAGGCACGATTTTTAGTATGATAGTTTTC**G**TAGTCT**T**AAA..3'

**Fig. S4. Determination of TSS and 5' ends of processing products of *rpsO* mRNA.**
Primer extension was performed as described in *Materials and Methods* using total RNA prepared from DB104 and DB104 (Δ*rny::spec*) and primer SB3403. A Sanger sequencing reaction performed on pUC19 with primer SB1171 was used as reference to calculate the location of the 5' ends. Black dot, TSS, the arrow shows the direction of transcription. The 5' ends of RNase Y processing products (PP1 and PP2) are underlined and in bold.

Figure S4 Ul Haq *et al.*

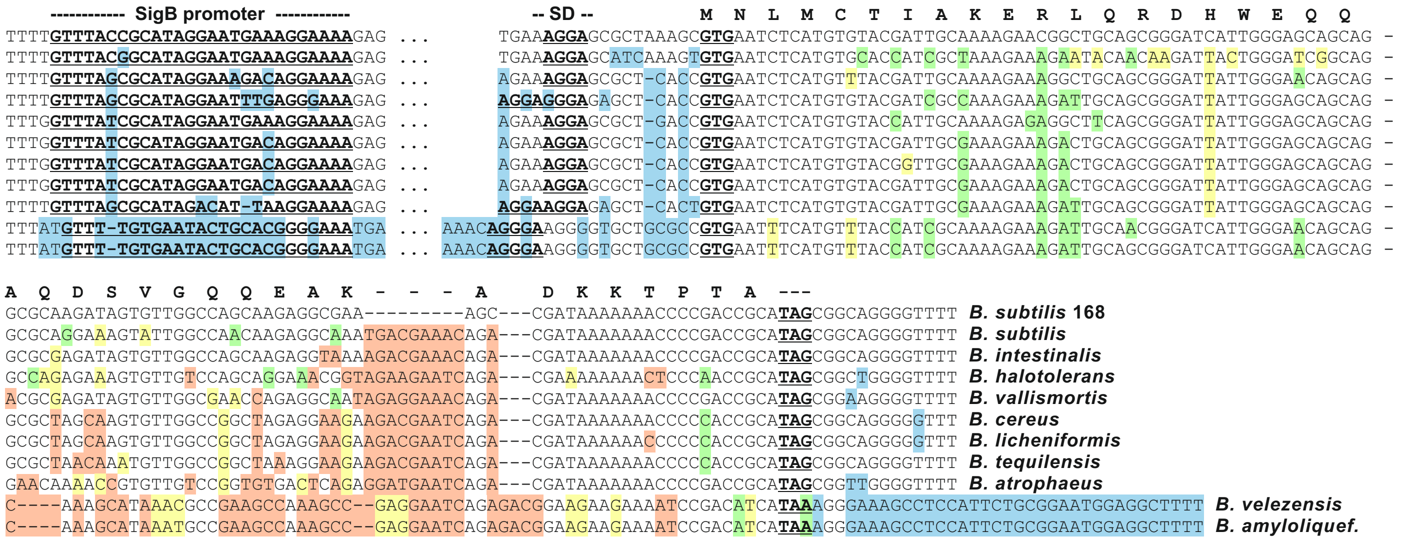


**Fig. S5. Gene conservation of *sr7p* homologues**

A BLAST search revealed homologous nucleotide sequences in 10 different species. Sequences were aligned to best matching the sequence of *B. subtilis* 168. In all cases, SigB-dependent promoters could be identified directly upstream. All genes comprise a rho-independent terminator. The putative promoter and SD sequences as well as start and stop codon of the ORF are marked in bold and underlined letters. The encoded aa sequence of *B. subtilis* 168 SR7P is shown above. A 38 to 44 nt region between promoter and SD sequence is not depicted (...). Sequence differences relative to the *sr7* gene from *B. subtilis* 168 are shaded in colour: blue, differences in non-coding regions; green, alterations not changing the encoded aa; yellow, alterations changing the encoded aa to a similar aa; orange, alterations changing the encoded aa to an non-similar aa. Sequence differences in the 5' part of the ORF result less frequently in an aa exchange, suggesting a higher aa conservation in this region.

**Table S1 Oligodeoxyribonucleotides used in this study**

| Oligo | Sequence | Purpose |
|---|---|---|
| SB2816 | 5'GATCCCCATGGTTGAAATCCCCTCAAAAACCCGATATAATGGGTTT AACCACCTTTTAAATGAAAGGAGCGCTAAAGCGTGAATCTCG | Fw translational *sr7p-lacZ* fusion |
| SB2817 | 5'AATTCGAGATTCACGCTTTAGCGCTCCTTTCATTTAAAAGGTGGTTA AACCCATTATATCGGGTTTTTGAGGGGATTTCAACCATGGG | Rev. translational *sr7p-lacZ* fusion |
| SB2843 | 5'GATAAGCTTGCAAAAAAAGACGTTTGCCTAAGGCAAACGTCTTTTT ACTTGTCATCGTCATCCTTGTA | C-FLAG elongation |
| SB2938 | 5'TATATTTATGTTACAGTAAATA | Fwd $Cm^R$ |
| SB2872 | 5'GATAAGCTTAGATAGGGCAAACACGACCAA | Fwd *pDRSR7P* |
| SB2878 | 5'CTTGTCATCGTCACCTTGTAATCGATATCATGATCTTTATAATCACC GTCATGGTCTTTGTAGTCTGCGGTCGGGGTTTTTTATCGGCTT | C-FLAG |
| SB2939 | 5'AAC TAA CGG GGC TAA GTG | Rev $Cm^R$ |
| SB2963 | 5'AGGATCTTTTTGTTTACCGCA | Fwd knock-ins |
| SB2964 | 5'CATGATGGATCCACCAGCGTCCAGATTTCTAG | Fwd front cassette |
| SB2966 | TATTACTGTAACATAAATATATGCGGTAAACAAAAAGATCCT | Rev front cassette |
| SB2967 | 5'CACTAACCTGCCCCGTTAGTTCAAGAGGCGAAAGCCGATAAA | Fwd rev cassette |
| SB2968 | 5'CATGATGAATCCTCGAGCATATGATAATGAAAG | Rev rev cassette |
| SB2978 | 5'CTTGTAATCGATATCATGATCTTTATAATCACCGTCATGGTCTTTGT AGTCTGCGGTCGGGGTTTTTTATCGGCTT | C-FLAG Knock-in |
| SB3013 | 5'AGCAGAAAACCCAGCTGTCATCGCGAA | Fwd front cassette |
| SB3014 | 5'TATTACTGTAACATAAATATACGGTAAACAAAA AGATCCTTTGCCT | Rev front cassette |
| SB3017 | CACTAACCTGCCCCGTTAGTTTCTTTTTGTTTACCGCATAGGAATGAA | Fwd 2360 insert/cat |
| SB3020 | 5'TCGAGCATATGATAATGAAAG | Rev back cassette |
| SB3039 | 5'TATATTTATGTTACAGTAAATAGAATTCTTGAAATCCCCTCAAAAAC | Fwd $spec^R$, $neo^R$ |
| SB3040 | 5'AACTAACGGGGCAGGTTAGTGATTACCAATTAGAATGAATATTTC | Rev $spec^R$ |
| SB3042 | 5'CACTAACCTGCCCCGTTAGTTAGATAGGGCAAACACGACCAA | Fwd 2360w/o pro |
| SB3156 | 5'ATGTTTTCATTTTGTTCCTTATCAATAAGGT | |
| SB3167 | 5'TATTACTGTAACATAAATATAGCAAAAAAAGACGTTTGCCTAAGGCA AACGTCTTTTTATTA | Rev $His_{10}$ elongation |
| SB3180 | 5'AACTAACGGGGCAGGTTAGTGTAATACTAGGAGAAGTTAATAAATA CGTAAC | Rev $neo^R$ |
| SB3182 | 5'GCCGTTCTTTTGCAATCGTACACATGAGATTCACGCT | Rev $neo^R$ |
| SB3226 | 5'TTTAAATGAAAGGAGCGCTAAAGCTAATAAAATCTCATGTGTACGA TTGCAAAAGAACGGCTGCAGCGGGAT | Fw first mutant, double stop codons |
| SB3227 | 5'ATCCCGCTGCAGCCGTTCTTTTGCAATCGTACACATGAGATTTTAT TAGCTTTAGCGCTCCTTTCATTTAAA | Rev first mutant |
| SB3243 | 5'CACTAACCTGCCCCGTTAGTTAAAGACGTTTGCCTTAGGCAAACGT CTTTTTTTGC | Fwd for 2360 ko |
| SB3267 | 5'TATTACTGTAACATAAATATATTATTACTTGTCATCGTCATCCTTGTA | Rev 1421, 3xFLAG |
| SB3269 | 5' AGGATGGAGATCCCTTTTCATTGTTTTTA | Fwd pfkA 3xFLAG |
| SB3270 | 5'CTTGTCATCGTCATCCTTGTAATCGATATCATGATCTTTATATCACC GTCATGGTCTTTGTAGTCGATAGACAGTTCTTTTGAAAGCTGATA | Rev pfkA 3xFLAG |
| SB3271 | CACTAACCTGCCCCGTTAGTTTGTACAGCTGAAGGCTGAAGATTTCA | Fwd pfkA back cass |
| SB3272 | 5' ACTTCCGGCAGCAGTTTCACCAGAAAGCAT | Rev pfkA back cass |
| SB3347 | 5'CCTAAGGCAAACGTCTTTTTATTAATGGTGGTGATGGTGATGATGG ATAGACAGTTCTTTTGAAGCTGATA | Rev pfkA-His6 6 His, 2 stops, BsrF |
| SB3439 | 5'CCTAAGGCAAACGTCTTTTTATTAATGGTGGTGATGGTGTGATGTG CGGTCGGGGTTTTTTTATCGGC | Rev SR7P-$His_6$ 2 stops, for pDR111 |
| SB3575 | 5'TATTACTGTAACATAAATATATTAGTGGTGATGATGGTGATGGTGG TGATGGTGTGCGGTCGGGGTTTTTTATCGGCTT | Rev SR7P $His_{10}$ Stop, overhang cat |
| SB3403 | 5' TGAGTTGGTTTTTACGCTCTTGAG | *rpsO* 5' mapping |

Sequencing primers

| | | |
|---|---|---|
| SB1171 | 5'GGATAACAATTTCACACAGGA | Rev primer for pUC19 |
| SB2976 | 5'AGGCTTACTTGTCTGCTTTCT | Rev primer for knock-ins/outs |
| SB2977 | 5'AATATGAGATAATGCCGACTG | Fwd primer for knock-ins/outs |
| SB3183 | 5'GTGGATGTGTCAAAACGCATACC | Knock-ins |
| SB3262 | 5'CCTATTTTATATCCATAGTTG | Knock-ins |

Oligos for PCR templates for *in vitro* transcription

| Name | Sequence | Purpose |
|---|---|---|
| SB2796 | 5'GAAATTAATACGACTCACTATAGGCTATGCGGTCGGGGTTTTTTTATC | Fwd oligo SR7 |
| SB2997 | 5'GGAGCGCTAAAGCGTGAATCT | Rev oligo SR7 |
| SB3169 | 5'AGGTGAAACAGGATGGCAATTA | Fwd oligo *rpsO* |
| SB3170 | 5'GAAATTAATACGACTCACTATAGGAGCCTAGTTTGTTAATTAACT | Rev oligo *rpsO* |
| SB3138 | 5'GAAATTAATACGACTCACTATAGGTGGAGACGGTGGGAGTCGAAC | Fwd oligo tmRNA |
| SB3139 | 5'TAAACACGCACTTAAATATAA | Rev oligo tmRNA |
| SB767 | 5'GGGTGTGACCTCTTCGCTATCGCCACC | 5S rRNA |

Oligos for 5' labelled RNase Y target RNAs

| | | |
|---|---|---|
| SB3404 | 5'GAAATTAATACGACTCACTATAGGGAAAGGGAAAAACGCCAGATATC | Fwd oligo *yitJ* |
| SB3405 | 5'ACTTTGTCAGTGATTTTGTCTCTT | Rev oligo *yitJ* |
| SB3406 | 5'GAAATTAATACGACTCACTATAGGGTAGTCTTAAAACCATTGCTT | Fwd oligo *rpsO* |
| SB3386 | 5'AAAAGCGGGAATCCTCCCGCT | Rev oligo *rpsO* |

**Table S2: Plasmids used in this study**

| Plasmid | Description | Reference |
|---|---|---|
| pUC19 | *E. coli* cloning vector, $Amp^R$, MCS | Sambrook *et al*. 1989 |
| pUC19-SR7P$_{Pro}$ | pUC19 with insert encoding the SR7P promoter with upstream sequence | This study |
| pDR111 | *E. coli* vector for insertion into the *B. subtilis amyE* locus, IPTG-inducible Hyperspank promoter, $Amp^R$, $Spec^R$ | van Ooij and Losick (2003) |
| pDRSR7P | pDR111 with promoterless *sr7p* gene | This study |
| pGAB1 | *E. coli* vector for insertion of *lacZ*-translational fusions into *B. subtilis amyE* locus, $Amp^R$, $Km^R$ | Müller *et al.,* 2019 |
| pGAB2360 | pGAB1 with translational *sr7p-lacZ* fusion under control of pIII | This study |
